# BRCA1-BARD1 combines multiple chromatin recognition modules to bridge nascent nucleosomes

**DOI:** 10.1101/2023.03.28.533771

**Authors:** Hayden Burdett, Martina Foglizzo, Laura J. Musgrove, Dhananjay Kumar, Gillian Clifford, Lisa J. Campbell, George R. Heath, Elton Zeqiraj, Marcus D. Wilson

## Abstract

Chromatin association of the BRCA1-BARD1 heterodimer is critical to promote homologous recombination repair of DNA double-strand breaks (DSBs) in S/G2. How the BRCA1-BARD1 complex interacts with chromatin that contains both damage induced histone H2A ubiquitin and inhibitory H4AK20 methylation is not fully understood. We characterised BRCA1-BARD1 binding and enzymatic activity to an array of mono- and di-nucleosome substrates using biochemical, structural, and single molecule imaging approaches. We find that the BRCA1-BARD1 complex preferentially interacts and modifies di-nucleosomes over mono-nucleosomes, allowing integration of H2A Lys-15 ubiquitylation signals with other chromatin modifications and features. Using high speed-AFM to provide real-time visualization of BRCA1-BARD1 complex recognising chromatin, we show a highly dynamic complex that bridges two nucleosomes and associates with the DNA linker region. Bridging is aided by multivalent cross-nucleosome interactions that enhance BRCA1-BARD1 E3 ubiqiutin ligase catalytic activity. Multivalent interactions across nucleosomes explains how BRCA1-BARD1 can recognize chromatin that retains partial di-methylation at H4 Lys-20 (H4K20me2), a parental histone mark that blocks BRCA1-BARD1 interaction with nucleosomes, to promote its enzymatic and DNA repair activities.

## Introduction

DNA double strand breaks (DSBs) are highly deleterious lesions that can form as a consequence of replication errors, chemical insults, or from the absorption of ionizing radiation (1). Cells have evolved several distinct and highly controlled DSBs repair pathways that are spatiotemporally regulated during the cell cycle to ensure the correct course of repair.

Chromatin profoundly shapes the response to DNA damage. The basic unit of chromatin is the nucleosome, comprising an octameric assembly of the four core histones which wraps and compacts roughly 147 bp of DNA (2,3). Nucleosomes form in a repeated pattern in the genome, with stretches of linker DNA separating the compacted nucleosome particles. Concurrent with their role as a structural scaffold, nucleosomes provide a landing platform to control chromatin processes that undergo a multitude of post-translational modifications (PTMs) (4). A host of these signalling events are induced upon DNA breakage (5). These PTMs either alter the local chemistry, affecting chromatin structure directly, or function to recruit effector proteins that multivalently recognise specific modifications, DNA and/or the nucleosome surface itself, thereby ensuring efficient signalling and repairing of DSBs.

Repair of DSBs can be roughly divided into two separate mechanisms, termed non-homologous end-joining (NHEJ) and homology directed repair (HDR). The choice between NHEJ and HDR is in part mediated by p53-binding protein 1 (53BP1) and breast cancer type 1 susceptibility protein (BRCA1). The 53BP1/RIF1/SHIELDIN complex promotes NHEJ repair (6,7), with 53BP1 preventing BRCA1-BARD1 recruitment during G1 phase of the cell cycle (8,9). In S/G2 phase, BRCA1 and DNA resection factors antagonize the 53BP1/RIF1/SHIELDIN complex and promote HDR (10,11). The mutual antagonism of 53BP1 and BRCA1 is a major driver of DSBs repair pathway choice.

BRCA1 is a tumour suppressor commonly mutated in a number of human malignancies, including breast and ovarian cancers (12–16). BRCA1 operates multiple steps of HDR-mediated DSBs repair and cell cycle checkpoint control by forming multiple protein complexes via intreraction with its Really Interesting New Gene (RING) domain, a coiled-coil region and two conserved BRCA1 C-terminal (BRCT) repeats. The BRCA1 RING partners with the analogous region in BARD1 (17) to form an obligate heterodimer. This is an E3 ubiquitin ligase, with specificity for Lys-125/127/129 on histone 2A (H2A) (18–20). Recent structural studies have shown that the BRCA1-BARD1 RING domains directly interact with the nucleosome surface and position the E2 ubiquitin-conjugating enzyme in proximity to H2A target lysine residues, thus providing important structural understanding for BRCA1-BARD1-mediated ubiquitin transfer and E3 ligase specificity (21,22). Recruitment of BRCA1 to sites of DNA damage is multimodal and is also mediated by additional interacting partners. (23–30). While 53BP1 and BRCA1-BARD1 are antagonistic and promote different repair pathways, both lie downstream of the enzymatic activity of E3 ligases RNF8 and RNF168 (31). Recent studies, including our own, have mapped the initial damage-induced recruitment of the BRCA1-BARD1 complex to chromatin through recognition of both induced DNA damage marks (H2AK15ub) and resident unmodified histone H4 at lysine 20 (H4K20me0). The BARD1 Ankyrin Repaet Domain (ARD) and BRCT modules are direct multivalent readers of these histone PTMs on nucleosomes, thus forming a second nucleosome-interacting module (in addition to the already described BRCA1-BARD1 RINGs) within the BRCA1-BARD1 complex (21,22,32–35).

These observations partially explain how cell cycle dependent DSBs pathway choice is centred on chromatin, and how recruitment of BRCA1-BARD1 is mediated via direct recognition of the absence of H4 Lys-20 methylation (32). Both 53BP1 and BRCA1-BARD1 recognise H2AK15ub. H4K20 di-methylation (H4K20me2) is a cell cycle stage-specific PTM, with high levels in G1 phase that are diluted semi-conservatively during DNA replication. H4K20me2 is selectively recognized by 53BP1, and thus represents a chromatin mark that specifically guides DNA repair pathway choice towards NHEJ. The BRCA1-BARD1 specificity for unmethylated H4K20, instead, marks freshly deposited post-replicative chromatin thereby promoting BRCA1-BARD1 retention on DNA damaged sites and HDR. Despite a number of recent studies explaining how minimal BRCA1-BARD1 domains interact with mono- nucleosomes, it is still unclear how larger BRCA1-BARD1 assemblies that contain the full complement of nucleosome binding domains recognise and modify chromatin, particularly in the context of both cognate recognition and inhibitory modifications.

Here we show that the fully assembled BRCA1-BARD1 complex integrates multiple chromatin binding regions to ensure recruitment to post- replicative chromatin. We have generated a variety of BRCA1-BARD1 fragments, including nearly full-length complexes, and designed specific mono- and di-nucleosomes to determine the exact substrate-binding preference of BRCA1-BARD1. Using a combination of biochemical, biophysical and structural biology approaches, we demonstrate that BRCA1-BARD1 is a specific reader of RNF8/RNF168-catalysed DNA damage marks and recognises lysine 63 (K63)-linked di-ubiquitin on H2A. The BRCA1-BARD1 RINGs and the BARD1 ARD-BRCTs nucleosome-interacting regions can work in tandem to preferentially engage on two adjacent nucleosomes, allowing BRCA1-BARD1 to bridge between partially H4K20 methylated chromatin found in S/G2 phases of the cell cycle where HDR is permissive.

## Materials and Methods

### Generation of plasmid constructs

A list of all BRCA1/BARD1 expression constructs used in this study is provided in Table S3.

#### Cloning of UBA1, UbcH5c, Ubc13, Mms2 and ubiquitin

UBA1, UbcH5c, Ubc13, Mms2, ubiquitin, and ub^G76C^ constructs were a gift from Professor Frank Sicheri’s lab. K63R mutant was introduced to ubiquitin by site directed mutagenesis.

#### Cloning of BRCA1/BARD1 complexes

GST-BARD1^425-777^ (GST-BARD1^ARD-BRCTs^) and 6xHis-MBP-BARD1^425-777^ (MBP-BARD1^ARD-BRCTs^) were used previously (33). Fused 6xHis-MBP-BRCA1^1-100^-BARD1^26-777^ (BRCA1^RING^- BARD1) construct was created by first cloning BARD1^26-777^ into 6xHis-MBP vector by Ligation Independent Cloning (LIC) using pCDNA5-FRT-TO BARD1 (Dr Daniel Durocher’s lab) as template. A BRCA^1-100^ construct from Addgene (36) was subsequently introduced into the His-MBP-TEV-BARD1^26-777^ vector by Gibson assembly in frame between the TEV site and BARD1 (37) according to manufacturer’s instructions (New England BioLabs).

To generate constructs for insect cells expression, genes for human or cat BRCA1^ΔExon11^ and full-length (FL) BARD1 were cloned into a pFL vector (38). A Flag tag or a double (d)StrepII and monomeric ultra-stable (mu)GFP tags, each followed by a PreScission protease site, were engineered at the N-terminus of BRCA1^ΔExon11^. A single 6xHis tag, also followed by a PreScission site, was introduced at the BARD1 N-terminus.

#### Cloning of histone constructs

Histone plasmids were purchased from Addgene. Mutants and variants have been reported previously (33,39,40), generated by site directed mutagenesis.

#### Nucleosome DNA

DNA for reconstituting mono- and di-nucleosomes was designed based on the Widom-601 strong positioning sequence (41) with 15 bp linker arms added. Di-nucleosome DNA contained 8 bp linker-147 bp Widom-601-30 bp linker-147 bp Widom-603 strong positioning sequence- 8 bp linker. All DNA was synthesised with DNA overhangs for cloning using Gibson assembly into pUC57 for storage and as use for PCR template.

### Protein expression and purification

#### Expression and purification of hUBA1 (E1)

hUBA1 was expressed in BL21 (DE3) cells in Terrific Broth (TB) media overnight (O/N) at 18°C with 1 mM Isopropyl ß-D-1-thiogalctopyranoside (IPTG). Cells were harvested by centrifugation at 5,500 x g at 4°C for 15 minutes, and stored at -80°C until required. Cells were resuspended in lysis buffer (50 mM Tris pH 7.6, 150 mM NaCl, 20 mM imidazole, 5% (v/v) glycerol, 0.075% (v/v) β-mercaptoethanol, 1 mM benzamidine, 0.8 mM phenylmethylsulfonyl fluoride (PMSF) and 0.3 mg.mL^-1^ lysozyme). Cells were lysed by sonication, before bacterial cell debris was pelleted by centrifugation for 30 minutes at 30,000 x g at 4°C. Lysate was briefly sonicated again to shear remaining genomic DNA. Clarified lysate was filtered using a 0.45 μm filter, prior to application to a 5 mL HiTrap chelating column (Cytiva) pre-equilibrated with low salt buffer (50 mM Tris pH 7.6, 150 mM NaCl, 20 mM imidazole, 5% (v/v) glycerol, 0.075% (v/v) β-mercaptoethanol and 1 mM benzamidine). Column was then washed with 4 column volume (CV) of low salt buffer, then 4 CV high salt buffer (500 mM NaCl instead of 150 mM) and again with 4 CV low salt buffer. hUBA1 was eluted over a linear gradient of 20 CV using elution buffer (50 mM Tris pH 7.6, 150 mM NaCl, 300 mM imidazole, 5% (v/v) glycerol, 0.075% (v/v) β-mercaptoethanol and 1 mM benzamidine). Fractions containing hUBA1 were combined into 3.5 kDa MWCO snakeskin and cleaved with TEV protease O/N at 4°C in 4 L of dialysis buffer (50 mM Tris pH 7.6, 150 mM NaCl, 20 mM imidazole, 5% (v/v) glycerol, 0.075% (v/v) β-mercaptoethanol and 1 mM benzamidine). The next day TEV protease, uncleaved hUBA1 and free 6xHis-tag was removed by immobilised metal affinity chromatography (IMAC), using the above low salt and elution buffers and after concentration applied to a HiLoad Superdex S200 16/600 pre-equilibrated with gel filtration buffer (50 mM Tris pH 7.5, 150 mM NaCl, 5% (v/v) glycerol and 1 mM tris(2-carboxyethyl)phosphine (TCEP). Fractions containing hUBA1 were pooled, concentrated using a 30 kDa MCWO centrifugal filter, aliquoted and snap frozen in liquid nitrogen before being stored at -80°C.

#### Expression and purification of UbcH5c

UbcH5c was expressed in BL21 (DE3) cells in 2xYeast Extract Tryptone (YT) media O/N at 18°C with 0.5 mM IPTG. Cells were harvested by centrifugation at 5,500 x g at 4°C for 15 minutes, and stored at -80°C until required. Cells were resuspended in lysis buffer (50 mM HEPES pH 7.5, 150 mM NaCl, 10 mM imidazole, 1 mM DTT) containing 1 mM 4-(2- aminoethyl)benzenesulfonyl fluoride hydrochloride (AEBSF), 1X protease inhibitor cocktail (2.2 mM PMSF, 2 mM benzamidine HCl, 2 μM leupeptin, 1 μg.mL^-1^ pepstainA), 4 mM MgCl_2_, 5 μg.mL^-1^ DNAse and 500 μg.mL^-1^ lysozyme. Cells were nutated at 4°C for 30-60 minutes, before additional lysis using a sonicator (2 s on, 2 s off for total 20 s at 50% amplitude). Bacterial cell debris was pelleted by centrifugation for 25 minutes at 39,000 x g at 4°C. Clarified lysate was filtered through a 0.4 μm filter, before being loaded onto a 5 mL HisTrap column pre-equilibrated with IMAC buffer A (50 mM HEPES pH 7.5, 150 mM NaCl, 10 mM imidazole, 1 mM dithiothreitol (DTT)). After extensive washing with IMAC buffer A, UbcH5c was eluted from the column using IMAC buffer B (50 mM HEPES pH 7.5, 150 mM NaCl, 500 mM imidazole, 1 mM DTT). Fractions containing UbcH5c were then concentrated using a 10 kDa MWCO centrifugal filter unit and further purified by size exclusion chromatography using a HiLoad Superdex S75 16/600 (Cytiva) pre-equilibrated with gel filtration buffer (10 mM HEPES pH 7.5, 150 mM NaCl, 5% (v/v) glycerol and 1 mM DTT). Fractions containing UbcH5c were pooled, concentrated using a 10 kDa MWCO centrifugal filter, aliquoted and snap frozen in liquid nitrogen before being stored at -80°C.

#### Expression and purification of Ubc13 and Mms2

Both Mms2 and Ubc13 were expressed in BL21 (DE3) cells in TB media O/N at 18°C with 1 mM IPTG. Cells were harvested by centrifugation at 5,500 x g at 4°C for 15 minutes, and stored at -80°C until required. Cells were resuspended in lysis buffer (50 mM Tris pH 7.6, 300 mM NaCl, 1 mM benzamidine, 0.8 mM PMSF and 0.3 mg.mL^-1^), and lysed by sonication. After sonication, 0.02% (v/v) and 10 mM MgCl_2_ was added to cell lysate, and incubated for 20 minutes at 4°C with rotation. Bacterial cell debris was pelleted by centrifugation for 30 minutes at 30,000 x g at 4°C, and lysate was briefly sonicated again to shear remaining genomic DNA. For Ubc13, clarified lysate was filtered using a 0.45 μm filter, prior to incubation with 2 mL glutathione (GSH) beads (pre-equilibrated in 1xPBS) for 2 h at 4°C with rotation. After incubation, beads were washed with 40 mL 1xPBS, 40 mL 1xPBS with 0.5% (v/v) Triton X- 100, followed by another 40 mL wash with 1xPBS. After last wash, GSH beads were resuspended in 2 mL 1x PBS, and incubated with Thrombin protease and 2 mM CaCl_2_ O/N at 4°C with rotation. After O/N incubation, beads were incubated for a further 1 h at room temperature prior to tubes being pelleted. Supernatant containing cleaved Ubc13 was further purified using a HiLoad Superdex S75 16/600 (Cytiva) pre-equilibrated with gel filtration buffer (50 mM Tris pH 7.5, 150 mM NaCl, 5% (v/v) glycerol and 1 mM DTT). Fractions containing Ubc13 were pooled, concentrated using a 3 kDa MCWO centrifugal filter, aliquoted and snap frozen in liquid nitrogen before being stored at -80°C. For Mms2, clarified lysate was applied to a 5 mL HiTrap chelating column (Cytiva) pre-equilibrated with low salt buffer (50 mM Tris pH 7.6, 350 mM NaCl, 20 mM imidazole, 5% (v/v) glycerol, 0.075% (v/v) β-mercaptoethanol and 1 mM benzamidine). Column was then washed with 4 CV of low salt buffer, then 4 CV high salt buffer (500 mM NaCl instead of 300 mM) and again with 4 CV low salt buffer. Mms2 was eluted over a linear gradient of 20 CV using elution buffer (50 mM Tris pH 7.6, 300 mM NaCl, 300 mM imidazole, 5% (v/v) glycerol, 0.075% (v/v) β-mercaptoethanol and 1 mM benzamidine). Fractions containing Mms2 were combined into 3.5 kDa MWCO snakeskin and cleaved with TEV protease O/N at 4°C in 4 L of dialysis buffer (50 mM Tris pH 7.6, 150 mM NaCl, 20 mM imidazole, 5% (v/v) glycerol, 0.075% (v/v) β-mercaptoethanol and 1 mM benzamidine). The next day TEV protease, uncleaved Mms2 and free 6xHis-tag was removed by IMAC, using the above low salt and elution buffers, and after concentration applied to a HiLoad Superdex S75 16/600 (Cytiva) pre-equilibrated with gel filtration buffer (50 mM Tris pH 7.5, 150 mM NaCl, 5% (v/v) glycerol and 1 mM DTT). Fractions containing Mms2 were pooled, concentrated using a 3 kDa MWCO centrifugal filter, aliquoted and snap frozen in liquid nitrogen before being stored at -80°C.

#### Expression and purification of ubiquitin variants

6xHis-Ubiquitin^G76C^ and 6xHis-Ubiquitin^K63R^ were expressed in BL21 (DE3) cells in 2xYT O/N at 16°C with 0.4 μM IPTG. Cells were harvested for 15 min at 5,500 x g at 4°C, and stored at -80°C until required. Cells were resuspended in lysis buffer (10 mM Tris pH 7.5, 500 mM NaCl, 10% (v/v) glycerol, 0.1% (v/v) Triton, 5 mM β-mercaptoethanol) containing 1 mM AEBSF, 1X protease inhibitor cocktail (2.2 mM PMSF, 2 mM benzamidine HCl, 2 μM leupeptin, 1 μg.mL^-1^ pepstainA), 4 mM MgCl2, 5 μg.mL^-1^ DNAse and 500 μg.mL^-1^ lysozyme. Cells were nutated at 4°C for 30-60 minutes, before additional lysis using a sonicator (2 s on, 2 s off for total 20 s at 50% amplitude). 15 mM imidazole was added to the lysate, prior to bacterial cell debris pelleting by centrifugation for 25 minutes at 39,000 x g at 4°C. Clarified lysate was filtered through a 0.4 μm filter, before being loaded onto a pre-equilibrated 5 mL HiTrap chelating column (Cytiva) with IMAC buffer A (20 mM Tris pH 7.5, 400 mM NaCl, 15 mM imidazole, 5 mM β-mercaptoethanol). Column was washed with 10 CV of IMAC buffer A, and eluted with a gradient over 12 CV of IMAC buffer B (20 mM Tris pH 7.5, 400 mM NaCl, 400 mM imidazole, 5 mM β-mercaptoethanol). Peak fractions were pooled and further purified by size exclusion chromatography on a HiLoad Superdex S75 16/600 (Cytiva) in a buffer containing 20 mM Tris pH 7.5, 150 mM NaCl and 5 mM DTT, to remove minor impurities and to ensure monodispersity. 5 mL of the pooled IMAC fractions was injected per run, and 6xHis-Ubiquitin containing peak fractions from SEC were pooled and dialysed extensively into 4 L of 1 mM acetic acid before being flash frozen in liquid nitrogen, lyophilised dried and stored at -20°C. *Expression and purification of BRCA1/BARD1 complexes from bacteria:*

GST-BARD1^ARD-BRCTs^ and 6xHis-MBP-BARD1^ARD-BRCTs^ were purified as described in Becker et al 2021. Constructs were expressed in BL21 (DE3 RIL) *E. coli* cells O/N with 200 μM IPTG at 16°C in 2xYT broth. Cell pellets were resuspended in lysis buffer (25 mM Tris pH 8.0, 300 mM NaCl, 0.1 % (v/v) Triton, 10% (v/v) glycerol, 5 mM β-mercaptoethanol, 1X protease inhibitor cocktail (2.2 mM PMSF, 2 mM benzamidine HCl, 2 μM leupeptin, 1 μg.mL^-1^ pepstainA), 4 mM MgCl_2_, 5 μg.mL^-1^ DNAse and 500 μg.mL^-1^ lysozyme) and nutated at 4°C for 30-60 minutes, before additional lysis using a sonicator (2 s on, 2 s off for total 20 s at 50% amplitude). Bacterial cell debris was pelleted by centrifugation for 25 minutes at 39,000 x g at 4°C and clarified lysate was filtered through a 0.4 μm filter. GST-BARD1^ARD-BRCTs^ clarified lysate was applied to a GSTrap HP column (Cytiva), extensively washed with wash buffer (25 mM Tris pH 7.5, 300 mM NaCl, 10% (v/v) glycerol, 5 mM β-mercaptoethanol) and eluted using wash buffer containing 30 mM reduced glutathione. Fractions containing GST-BARD1^ARD-BRCTs^ were pooled and concentrated using a 30 kDa MWCO centrifugal filter. 6xHis-MBP-BARD1^ARD-BRCTs^ clarified lysate was supplemented with 15 mM imidazole before being applied to a chelating HP column pre-charged with nickel ions, extensively washed with wash buffer (25 mM Tris pH 7.5, 500 mM NaCl, 10% (v/v) glycerol, 15 mM imidazole, 2 mM β-mercaptoethanol) and eluted with elution buffer (25 mM Tris pH 7.5, 500 mM NaCl, 10% (v/v) glycerol, 300 mM imidazole, 2 mM β-mercaptoethanol) over a 15 CV gradient. Fractions containing 6xHis-MBP-BARD1^ARD-BRCTs^ were pooled and concentrated using a 30 kDa MWCO centrifugal filter. Both proteins were further purified by size exclusion chromatography using a HiLoad Superdex 200 16/600 (Cytiva) in gel filtration buffer (15 mM HEPES pH 7.5, 150 mM NaCl, 1 mM DTT, 5% (v/v) glycerol). Fractions were analysed by SDS-PAGE and those containing GST-BARD1^ARD-BRCTs^ or 6xHis-MBP-BARD1^ARD-BRCTs^ were pooled, concentrated in a 30 kDa MWCO centrifugal filter, and flash frozen in liquid nitrogen and stored at -80°C.

6xHis-MBP-BRCA1^RING^-BARD1 was expressed in BL21 (DE3 RIL) *E. coli* cells O/N with 500 μM IPTG at 16°C in 2xYT broth. Cell pellets were resuspended in lysis buffer (50 mM HEPES pH 7.5, 500 mM NaCl, 0.1 % (v/v) Triton, 10% (v/v) glycerol, 5 mM β-mercaptoethanol, 1X protease inhibitor cocktail (2.2 mM PMSF, 2 mM benzamidine HCl, 2 μM leupeptin, 1 μg.mL^-1^ pepstainA), 4 mM MgCl_2_, 5 μg.mL^-1^ DNAse and 500 μg.mL^-1^ lysozyme) and nutated at 4°C for 30-60 minutes, before additional lysis using a sonicator (2 s on, 2 s off for total 20 s at 50% amplitude). Bacterial cell debris was pelleted by centrifugation for 25 minutes at 39,000 x g at 4°C and clarified lysate was filtered through a 0.4 μm filter. 6xHis-MBP-BRCA1^RING^- BARD1 clarified lysate was supplemented with 10 mM imidazole before being applied to a HiTrap chelating HP column (Cytiva) pre-charged with nickel ions, extensively washed with wash buffer (50 mM HEPES pH 7.5, 500 mM NaCl, 10% (v/v) glycerol, 10 mM imidazole, 2 mM β-mercaptoethanol, 0.1 mM AEBSF) and eluted with elution buffer (50 mM HEPES pH 7.5, 500 mM NaCl, 10% (v/v) glycerol, 300 mM imidazole, 2 mM β-mercaptoethanol, 0.1 mM AEBSF) over a 15 CV gradient. Fractions containing 6xHis-MBP-BRCA1^RING^-BARD1 were pooled and diluted to 200 mM NaCl in SP buffer No Salt (10 mM HEPES pH 7.0, 1 mM DTT, 5% (v/v) glycerol, 0.1 mM AEBSF), before further purification by ion exchange chromatography. Diluted 6xHis-MBP-BRCA1^RING^-BARD1 fractions were applied to a pre-equilibrated SP HP column and washed extensively with SP A buffer (10 mM HEPES pH 7.0, 200 mM NaCl, 1 mM DTT, 5% (v/v) glycerol and 0.1 mM AEBSF), before eluting with SP B buffer (10 mM HEPES pH 7.0, 1000 mM NaCl, 1 mM DTT), across an 18 CV gradient. Fractions containing 6xHis-MBP-BRCA1^RING^-BARD1 were then pooled, concentrated in a 30 kDa MWCO centrifugal filter unit, before being applied to a HiLoad Superdex 200 16/600 (Cytiva) pre-equilibrated with gel filtration buffer (10 mM HEPES pH 7.5, 300 mM NaCl, 5% (v/v) glycerol, 1 M DTT, 2 μM ZnCl_2_ and 0.1 mM AEPSF). Fractions containing 6xHis-MBP-BRCA1^RING^-BARD1 were then pooled, concentrated, aliquoted and flash frozen in liquid nitrogen, before being stored at -80°C.

#### Expression and purification of BRCA1:BARD1 complexes from insect cells

To co-express BRCA1:BARD1 complexes from insect cells, bacmid DNA was generated in DH10MultiBac Turbo cells (ATG Biosynthetics) following manufacturer’s protocol, and virus amplification in *Spodopetera frugiperda 9* (*Sf9*) cells (ThermoFisher) was performed using standard procedures and as described in Zeqiraj et al. (42). For co-expression of the human and cat BRCA1:BARD1 variants, *Trichoplusia Ni* (*Tni*) cells (Oxford Expression Technologies) were infected with baculoviruses encoding each protein complex. Following 48 h after infection, cells were harvested by centrifugation at 500 g for 15 min and protein purification was carried out as outlined below.

Cells expressing human and cat Flag-BRCA1^ΔExon11^:6xHis-FL BARD1 were resuspended in 100 mL ice-cold low salt buffer [50 mM Tris-HCl pH 7.6, 300 mM NaCl, 20 mM Imidazole, 5% (v/v) glycerol, 0.075% (v/v) β-mercaptoethanol and 1 mM benzamidine] supplemented with one tablet of Pierce protease inhibitor cocktail (ThermoFisher Scientific) and lysed by sonication using a Sonics Vibracell instrument (1 s ON/3 s OFF, at 40% amplitude for 4 min). The cell lysates were cleared by centrifugation at 30,000 g for 30 min at 4°C, and the soluble fractions were sonicated (1 s ON/3 s OFF, at 40% amplitude for 2 min) and subsequently passed through a 0.45 μm filter (ThermoFisher Scientific). Filtered lysates were loaded onto a 5 mL HisTrap HP column (GE Healthcare), which was washed with four CV of low salt buffer, four CV of high salt buffer (low salt buffer containing 500 mM NaCl), and four CV of low salt buffer. A linear 20 CV gradient, from 20 mM to 300 mM Imidazole, was used to elute each complex. Eluted peaks were analyzed by SDS-PAGE, and fractions containing human or cat complexes were pooled and dialyzed for 4 h against 4 L dialysis buffer [25 mM HEPES pH 7.5, 500 mM NaCl, 5% (v/v) glycerol, 0.075% (v/v) β-mercaptoethanol and 1 mM benzamidine] at 4°C. The dialyzed samples were subsequently incubated with 1 mL Pierce Flag-affinity resin (ThermoFisher Scientific), pre-equilibrated in dialysis buffer, for 2 h at 4°C in rotation. After washing the resins two times with 50 mL dialysis buffer and twice with 50 mL wash buffer [dialysis buffer containing 300 mM NaCl], human and cat complexes were eluted by subsequent washes (1 mL each) with elution buffer [wash buffer supplemented with 100 μg.mL^-1^ Flag peptide (Pierce)]. Fractions containing human or cat BRCA1^ΔExon11^:FL BARD1 were pooled, diluted three times in 25 mM HEPES pH 7.5, 5% (v/v) glycerol and 1 mM TCEP, and loaded onto a 5 mL HiTrap SPHP column (GE Healthcare). A linear 5 CV gradient, from 0.1 M to 1 M NaCl, was used to elute each complex. Eluted peaks were analyzed by SDS-PAGE, and fractions containing >95% pure human or cat proteins were combined, concentrated to 2-4 mg.mL^-1^, snap-frozen in liquid nitrogen and stored at -80°C.

Purifications of human and cat dStrepII-muGFP-BRCA1^ΔExon11^:6xHis-FL BARD1 complexes were carried out in a similar manner, with the exception that Strep-affinity chromatography was used as a second purification step in place of Flag-affinity. Briefly, fractions containing each protein complex after nickel-affinity chromatography were pooled and dialyzed for 4 h against 4 L binding buffer [50 mM Tris pH 7.5, 300 mM NaCl, 5% (v/v) glycerol and 1 mM TCEP] at 4°C. The dialyzed samples were subsequently loaded onto a 1 mL StrepTrap HP column (GE Healthcare), which was washed with 20 CV of binding buffer. A linear 15 CV gradient, from 0 mM to 5 mM desthiobiotin (IBA Lifesciences GmbH), was then used to elute each complex. Fractions containing human or cat complexes were pooled and subjected to ion-exchange chromatography as described above. Peaks containing >95% pure human or cat proteins were combined, concentrated to 4-6 mg.mL^-1^, snap-frozen in liquid nitrogen and stored at -80°C.

#### Expression and purification of histone proteins

Unmodified human histones, as well as various cysteine mutants, were expressed and purified as previously described (39,40,43). Briefly, histones were expressed in BL21 (DE3 RIL) cells and resolubilised from inclusion bodies. Histones were further purified by cation exchange chromatography prior to dialysis in 1mM acetic acid and lyophilisation.

Protein concentrations were determined via absorbance at 280 nm using a Nanodrop One spectrophotometer (Thermo Scientific) and comparisons/purity was assessed by subsequent SDS-PAGE with comparison to known amounts of control proteins (Supplemental Figures S1A, S2A).

### Chemical modifications of histones

#### H2A and H2B ubiquitylation

Crosslinking reactions were performed as previously described (39,44) with some modifications to improve yield. Lyophilised 6xHis-ubiquitin^G76C^ and H2A^K15C^ were resuspended in 10 mM acetic acid at 80 mg.mL^-1^, before being mixed together to a final concertation of 20 mg.mL^-1^ in 10 mM acetic acid. The reaction was setup as follows, 1,3-dibromoacetone (DBA), freshly diluted in N,N’-dimethyl-formamide (DMF) was added to a buffered Tris solution (100 mM Tris pH 7.5) to a final concentration of 4.2 mM and mixed thoroughly. The 6xHis- ubiquitin^G76C^ and H2A^K15C^ mixture was mixed thoroughly prior to addition to the buffered DBA, and mixed thoroughly by pipetting. This modification ensured cysteine resides were kept protonated in low pH prior to mixing with buffer reactants. The reaction was left to proceed for 30-60 minutes on ice, before being quenched by addition of 20 mM β-mercaptoethanol. The reaction mixtures were then run on a SDS-PAGE gel to determine efficiency, then subjected to purification.

Ubiquitylated histones were purified in two steps. First, the reaction was purified by ion exchange chromatography, to remove the majority of unreacted ubiquitin and di-ubiquitin, and a further IMAC purification step to remove unreacted H2A and H2A dimers. After quenching, the crosslinking reaction was diluted 1:5 into SP A buffer (20 mM Tris pH 7.5, 50 mM NaCl, 4 mM β-mercaptoethanol, 7 M urea) and pH was adjusted to 7.5, prior to loading onto a pre-equilibrated HiTrap SP HP column (Cytiva). After extensive washing with SP A buffer, unreacted histone, di-histone and ubiquitylated histones were eluted with a gradient over 18 CV of 0-70% SP buffer B (20 mM Tris pH 8.8, 900 mM NaCl, 4 mM β-mercaptoethanol, 7 M urea). Fractions containing ubiquitylated histones were pooled and subjected to further purification by IMAC. 15 mM imidazole and 250 mM NaCl were added to pooled SP fractions, and samples was loaded onto a pre-equilibrated IMAC column (20 mM Tris pH 7.5, 400 mM NaCl, 15 mM imidazole, 5 mM β-mercaptoethanol, 5 M urea). After extensive washing with 5% IMAC buffer B (20 mM Tris pH 7.5, 400 mM NaCl 400 mM imidazole, 5 mM β- mercaptoethanol, 5 M urea), ubiquitylated histones were eluted from IMAC column using 100% IMAC buffer B. Peak fractions containing ubiquitylated histones were pooled, extensively dialysed in 1 mM β-mercaptoethanol, and lyophilised dried. The 6xHis-tag was removed from the ubiquitylated histone by cleavage with TEV protease. The lyophilised ubiquitylated histones were rehydrated in TEV buffer (20 mM Tris pH 7.5, 100 mM NaCl, 2 mM β-mercaptoethanol, 4 mM citrate, 1 M urea) and incubated O/N at 4°C with 1:200 ratio of TEV protease. The next day TEV protease, uncleaved ubiquitylated histones and free 6xHis- tag was removed by IMAC, using the above IMAC buffers. Cleaved ubiquitylated histones were then dialysed into 1 mM β-mercaptoethanol, lyophilised dried and stored at -20°C.

#### Ubiquitin chain assembly on H2A-Kc15ub

To generate K63-linked di-ubiquitin chains on H2A, 6xHis-ubiquitin^K63R^ and H2AKc15ub was used in a ubiquitylation reaction containing 50 mM Tris pH 7.6, 150 mM NaCl, 2.5 mM ATP, 5 mM MgCl_2_, 1 mM β-mercaptoethanol, 140 μM 6xHis-ubiquitin^K63R^, 20 μM H2AKc15ub, 600 nM E1, 8 μM Ubc13, 8 μM Mms2, 10 mM creatine phosphate and 0.6 U.mL^-1^ creatine phosphokinase. After 2 h at 37°C with shaking at 200 rpm, di-ubiquitylated histones were purified from reaction components using the same purification strategy as making mono-ubiquitylated histones by chemical crosslinking. The 6xHis tag on ubiquitin^K63R^ was cleaved as above, and cleaved ubiquitylated histones were then dialysed into 1 mM β- mercaptoethanol, lyophilised dried and stored at -20°C.

#### H4 methyl lysine analogue preparation

H4K20C was expressed and purified as described for other histones. The H4K20C protein was alkylated essentially as previously described (33,45). Briefly H4K20C was resuspended in denaturing buffer and (2-chloroethyl)-dimethylammonium chloride reagent was added and incubated at 20°C for 2 h. Reaction was quenched with β-mercaptoethanol and desalted using PD-10 columns (GE healthcare). Extent of reaction was checked using 1D intact weight ESI mass spectrometry (SIRCAMs, School of Chemistry, University of Edinburgh).

### Reconstitution of histone octamers

Histone octamers containing the purified ubiquitylated histones were reconstituted as preciously described by (39,43). Histones were rehydrated to ∼5 mg.mL^-1^ in unfolding buffer (20 mM Tris pH 7.5, 7 M Guanidine-HCl, 10 mM DTT) for 30 minutes at room temperature. Unfolded histones were mixed together in a 1.2:1.2:1:1 molar ratio of H2Aub:H2B:H3:H4 and diluted in unfolding buffer to a final concentration of 2 mg.mL^-1^. The histone mixture was then dialysed extensively in refolding buffer (15 mM Tris pH 7.5, 2 M NaCl, 1 mM EDTA, 5 mM β- mercaptoethanol) O/N at 4°C. Refolded octamers were separated from soluble aggregates, tetramers, dimers and unpaired histones by size exclusion chromatography on a Superdex S200 16/60 (Cytiva) in refolding buffer. Peak fractions corresponding to the refolded octamers were pooled, concentrated and then used immediately for nucleosome reconstitution, or stored at -20°C after addition of 50% (v/v) glycerol.

Octamers for asymmetric nucleosomes were formed and purified as described by Li and Shogren-Knaak (46) with some modifications. H2A, H2AKc15ub-6xHis, H2B, H3 and H4 were combined together in 1.17:0.13:1.3:1:1 molar ratio in unfolding buffer (20 mM Tris pH 7.5, 2 M NaCl, 5 mM DTT), before being dialysed into refolding buffer (15 mM Tris pH 7.5, 2 M NaCl, 1 mM EDTA, 5 mM β-mercaptoethanol). Octamers were subjected to IMAC to select for asymmetric and H2AKc15ub-His containing octamers. Octamers were applied to a 1 mL HisTrap column (Cytiva) pre-equilibrated with IMAC buffer A (15 mM Tris pH 7.5, 2 M NaCl, 10 mM imidazole, 5 mM β-mercaptoethanol), and washed extensively with IMAC buffer A, before elution with IMAC buffer B (15 mM Tris pH 7.5, 2 M NaCl, 300 mM imidazole, 5 mM β- mercaptoethanol). Octamers were then purified from soluble aggregates, and unpaired dimers by size exclusion chromatography using a Superdex S200 16/600 pre-equilibrated in refolding buffer (see above). Peak fractions were pooled and concentrated before being used to wrap nucleosomes, or stored at -20°C following the addition of glycerol, final concentration 50% (v/v).

### Nucleosome reconstitution

FAM labelled 175-bp Widom-601 and di-nucleosome DNA fragments for nucleosome reconstitution were generated by PCR amplification and purified as previously described (33,41). Fluorescent dyes were incorporated in the primers (IDT technologies). 384x100 μL reactions using Pfu polymerase and HPLC pure oligonucleotides were pooled, filtered through a 0.4 μm filter, and applied to a 6 mL ResourceQ column (Cytiva) pre-equilibrated with 10 mM Tris pH 7.5 and 1 mM EDTA. The column was then washed extensively with 500 mM NaCl, before eluting across a 12 CV gradient from 500 M NaCl to 900 mM NaCl. Fractions were analysed by native PAGE, and fractions containing the desired product were pooled, and concentrated by ethanol precipitation.

Nucleosomes were reconstituted as previously described (43), with some minor modifications. Purified octamers were incubated with DNA and wrapped using an 18 h exponential salt reduction gradient. The extent and purity of nucleosomes wrapping was checked by native PAGE and SDS-PAGE analysis (Supplementary Figures S1B, C, F and G, S3D and E, S4C and D). Free DNA was removed from mononucleosomes by partial PEG precipitation, using 9% (w/v) PEG 6000. Di-nucleosomes were reconstituted in the same manner as ‘mono’-nucleosomes, with the exception that a DNA:octamer ratio of 0.35-0.45:1 was used instead, and no PEG precipitation was performed.

To generate heterotypic di-nucleosomes, a 1:1 molar ratio of a tagged- and untagged- octamer was used and the tag position was dependent on the heterotypic di-nucleosomes to be made. When making the unmodified-H2AKc15ub di-nucleosomes, the H2AKc15ub containing octamer retained the 6xHis tag; when making the H2AKc15ub- H2AKc15ub/H4Kc20me2 di-nucleosomes, the H2AKc15ub/H4Kc20me2 containing octamer retained the 6xHis tag. To purify heterotypic di-nucleosomes, samples were incubated with 20 μL Ni-NTA beads pre-equilibrated with IMAC A buffer (15 mM HEPES pH 7.5, 150 mM NaCl, 5 mM imidazole, 10% (v/v) glycerol, 2 mM β-mercaptoethanol) for 2 h at 4°C with rotation. The unbound fraction was removed by transferring the resin to Pierce Micro-Spin columns (Thermo Fisher Scientific), and centrifugation at 100 x g for 15 s at 4°C. The resin was washed three times with 100 μL of IMAC A buffer, with centrifugation between each step as above. 6xHis- tagged nucleosomes were then eluted in 20 μL of IMAC A buffer with increasing increments of imidazole. To determine species of nucleosome present, both native PAGE and SDS-PAGE gels of purified nucleosomes were used. After purification, nucleosomes were buffer exchanged into 10 mM HEPES pH 7.5, 100 mM NaCl, 1 mM DTT, 0.1 mM AEBSF and used for ubiquitylation assays and EMSAs.

### Nucleosome pull-down assays

Pull downs were performed as previously described (33). Briefly, 8.5 μg GST-GST or 8.5 μg GST-BARD1 or 8.5 μg GST-BARD1 + 8.5 μg GST-GST was immobilised on 15 μL Glutathione Sepharose 4B beads (GE Healthcare) in 100 μL pulldown buffer (50 mM Tris pH 7.5, 150 mM NaCl, 0.02% (v/v) NP40, 0.1 mg.mL^-1^ BSA, 10% (v/v) glycerol, 1 mM EDTA, 2 mM β- mercaptoethanol) for 2 h at 4°C with rotation. Beads were washed by centrifugation at 500 x g for 1 minute at 4°C and resuspension in 1 mL of pulldown buffer. Beads were then incubated with 1.25 μg nucleosomes in the same buffer for 2 h at 4°C with rotation. Pulldowns were washed three times as above, before the beads were then resuspended in 2xSDS loading dye and boiled for 5 minutes to release bound proteins. 5% of input nucleosomes were loaded as a control alongside 3 μL of the pulldown reactions. Samples were analysed by western blotting for GST, H2A, H2B and H3.

### Electrophoretic mobility shift assays

6-Carboxyfluorescein (5’ 6-FAM) labelled nucleosomes (at 2 nM each) were incubated with various concentration ranges, as noted in figure legends of particular experiment, of 6xHis- MBP-BARD1^ARD-BRCTs^, 6xHis-MBP-BRCA1^RING^-BARD1 or dStrepII-muGFP-BRCA1^Δ11^:6xHis-FL BARD1 proteins in EMSA buffer (50 mM Tris pH 7.5, 150 mM NaCl, 0.02% (v/v) NP40, 0.1 mg.mL^-1^ BSA, 10% (v/v) glycerol, 1 mM DTT, 8% (w/v) sucrose, 0.01% (w/v) bromophenol blue) to a final volume of 12 μL. Samples were incubated on ice for 1 h, and products were separated by native-PAGE using 1xTris Glycine as running buffer for 90 minutes at 4°C. Gels were imaged for FAM signal (Excitation Blue light, Emission 532nm) using BioRAD ChemiDoc MP, and stained using Diamond DNA Stain (Promega).

### Microscale thermophoresis

Microscale thermophoresis (MST) measurements were performed on an NT.115 Monolith instrument (NanoTemper Technologies) using premium capillaries (NanoTemper; catalogue number: MO-K025). Reconstituted 5’ 6-FAM labelled nucleosome variants (at 20 nM each) were mixed with a dilution series of human 6xHis-MBP-BARD1^ARD-BRCTs^ or Flag- BRCA1^Δ11^:6xHis-FL BARD1 in a buffer containing 50 mM HEPES pH 7.5, 150 mM NaCl, 0.1 mg.mL^-1^ BSA, 10% (v/v) glycerol, 0.02% (v/v) NP40 and 1 mM DTT, and incubated at room temperature for 30 min. Dilution series ranged from 0.122 nM to 4000 nM (for 6xHis-MBP- BARD1^ARD-BRCTs^) and from 0.122 nM to 2000 nM (for Flag-BRCA1^Δ11^:6xHis-FL BARD1). Dilutions ranging between 4000 nM or 2000 nM-7.812 nM and between 7.812 nM-0.122 nM were performed in two-fold and four-fold steps respectively. All MST measurements were carried out using 50% LED power and 20% MST power, with 30 s laser ON and 5 s laser OFF time. MST data were analyzed in GraphPad Prism 7 v7.0c (GraphPad Software) using the thermophoresis and temperature jump data, and dissociation constant (*K*_d_) values were calculated by fitting the data with the GraphPad Prism built-in total binding equation for one site: y = Bmax*X/(*K*_d_ + X) + NS*X + Background. A summary of *K*_d_ values is shown in Supplementary Table S1.

### Nucleosome ubiquitylation assays

Assays were performed in a reaction buffer containing 50 mM Tris pH 7.5, 120 mM NaCl, 5 mM ATP, 5 mM MgCl_2_, 1 mM DTT, 8 μM Ubiquitin-Alexa647, 20 μM Ubiquitin, 500 nM UbcH5c, 50 nM E3 and 150 nM E1. Assays using mono-nucleosomes contained 1.2 μM nucleosome, whilst assays using di-nucleosomes contained 0.6 μM di-nucleosome (equal concentration of substrate H2A). Reactions were performed at 32°C with shaking at 700 rpm in a thermomixer (Eppendorf). 4 μL of reaction prior to addition of E1 served as a zero time point. After addition of E1, indicated time points were taken by removal of 4 μL of reaction and added to 2X SDS loading dye. Samples were run on SDS-PAGE gels (either 4-20% gradient or 17%) and imaged for Alexa647 fluorescence (excitation Red light, emission 700nm +/- 50nm) in a ChemiDoc instrument (Bio-Rad). Proteins were subsequently transferred and blotted, or stained with colloidal Coomassie to determine loading.

### Cryo-electron microscopy – grids preparation and data collection

*Cryo-EM grids preparation for BRCA1^ΔExon11^:FL BARD1 in complex with H2AKc15ub nucleosomes:* To obtain a cryo-EM structure of dStrepII-muGFP-BRCA1^ΔExon11^/6xHis-FL BARD1 bound to H2AKc15ub nucleosomes, cat (*Felis catus*) BRCA1^ΔExon11^/FL BARD1 (at 3 μM) was incubated with H2AKc15ub nucleosomes (at 1.5 μM) in 20 mM HEPES pH 7.5, 50 mM NaCl and 1 mM DTT for 1 h on ice. Quantifoil R3.5/1 200-mesh grids (Quantifoil Micro Tools GmbH) were glow-discharged for 30 s at 40 mA using a GloQube (Quorum) glow discharge unit. Cryo-EM grids were prepared by applying 3 μL of the BRCA1^ΔExon11^:FL BARD1 with H2AKc15ub nucleosomes mixture onto the glow-discharged Quantifoil grids, followed by immediate blotting (blot force = 0 N, blot time = 8 s) and plunge-freezing in liquid ethane cooled by liquid nitrogen, using a FEI Vitrobot IV (ThermoFisher) at 100% relative humidity and with a chamber temperature set at 4°C. A dataset was collected on a FEI Titan Krios transmission electron microscope (ThermoFisher) operating at 300 keV, using a magnification of 96,000x and a final calibrated object sampling of 0.86 Å/pixel. A total of 16,015 movies were recorded using the EPU automated acquisition software on a FEI Falcon IV direct electron detector in counting mode (47). A dose per physical pixel/s of 5.38 was used for each exposure movie, resulting in a total electron dose of 36.4 e^-^/Å^2^, fractionated across 172 EPU frames. These were then grouped into 26 frames, resulting in an electron dose of 0.8 e^-^/Å^2^ per frame. 17 exposures per hole were collected with a total exposure time of 5 s, and defocus values ranging from -1.7 μm to -3.1 μm. Detailed information on data collection and structure refinement is shown in Supplementary Table S2.

*Cryo-EM grids preparation for the isolated BRCA1^ΔExon11^:FL BARD1 complex:* UltrAuFoil R1.2/1.3 300-mesh gold grids (Quantifoil Micro Tools GmbH) were cleaned via indirect plasma using a Tergeo EM plasma cleaner (Pie Scientific) at 15 W for 1 min, and with flow rates for nitrogen, oxygen and argon gasses of 20.0, 19.8 and 29.0 sscm respectively. Cryo-EM grids were prepared by applying 3 μL of the purified cat Flag-BRCA1^ΔExon11^:6xHis-FL BARD1 complex (at 0.2 mg.mL^-1^) onto the glow-discharged gold grids, followed by immediate blotting (blot force = 6 N, blot time = 6 s) and plunge-freezing in liquid ethane cooled by liquid nitrogen, using a FEI Vitrobot IV (ThermoFisher) at 100% relative humidity and with a chamber temperature set at 4°C. A dataset was collected on a FEI Titan Krios transmission electron microscope (ThermoFisher) operating at 300 keV, using a total electron dose of 73 e^-^/Å^2^, a magnification of 75,000x, and a final calibrated object sampling of 1.065 Å/pixel. A total of 227 movies were recorded using the EPU automated acquisition software on a FEI Falcon III direct electron detector in integrating mode (47). Each exposure movie (one per hole) had a total exposure time of 1.7 s collected over 50 frames, with an electron dose of 1.46 e^-^/Å^2^ per frame and defocus values ranging from -1.5 μm to -3.0 μm. Detailed information on data collection is shown in Supplementary Table S2.

### Cryo-electron microscopy – data processing

*Processing of the BRCA1^ΔExon11^/FL BARD1 in complex with H2AKc15ub nucleosomes dataset:* A schematic of the data processing pipeline in shown in Supplementary Figure. S5, while further details on the reported map and model are available in Supplementary Table S2. Image processing was carried out using a combination of RELION v3.1.2 (48) and cryoSPARC v3.2.0 (49). Drift-corrected averages of each movie were created using RELION’s implementation of MotionCor2 (50), and real-time contrast transfer function (CTF) parameters of each determined using CTFFIND-4.1 (51). Both motion correction and CTF estimation were carried out on-the-fly (47). Initially, 2,113 particles were manually picked and used to train crYOLO v1.6.1 (52). This trained model was used for picking on 3,203 movies (20% of the collected dataset), resulting in 1,440,728 particles. Particles were imported in RELION, extracted using a box size of 320 pixels and a binning factor of two, and subsequently subjected to reference-free 2D classification with a mask diameter of 240 Å. After visual inspection, high quality 2D classes (983,454 particles) were selected and used as references for auto-picking. Following auto-picking, 7,204,662 particles were extracted as indicated above and subsequently imported in cryoSPARC for iterative rounds of reference-free 2D classification with a mask diameter of 240 Å. Following 2D classification, a total of 1,405,673 particles were retained and used to generate three initial 3D volumes. These models were subsequently subjected to 3D classification using heterogeneous refinement with C1 symmetry, which yielded a well-resolved map (953,766 particles) containing the isolated ankyrin repeat domain (ARD) and tandem BRCA1 C-terminal repeats (BRCTs) of BARD1 bound to H2AKc15ub nucleosomes. No additional densities belonging to either BARD1 or BRCA1^ΔExon11^ were visible in the model. This model was subsequently used as a template for non-uniform 3D refinement, generating a map with a global resolution of 3.55 Å. The 953,766 particles belonging to this model were then imported in RELION, and re-extracted using a box size of 320 pixels without binning. The corresponding map was also re-scaled to the same box and pixel sizes using “relion_image_handler”. The resulting particles and model were subsequently subjected to 3D refinement, per-particle CTF correction and particle polishing in RELION (48). Post-processing was used to appropriately mask the models and estimate and correct for the B-factor of the map while the Phenix auto-sharpening tool was employed to adjust the resolution dependence of the map and improve its clarity. The final resolution (3.40 Å) was determined using the gold-standard Fourier shell correlation criterion (FSC = 0.143). The local resolution of the reconstructed map was determined using the local resolution implementation in RELION. *Processing of the BRCA1^ΔExon11^:FL BARD1 dataset:* A schematic of the data processing pipeline in shown in Supplementary Figure. S7B, and details on the reported maps are available in Supplmentary Table S2. Image processing was carried out using a combination of RELION v3.1.2 (48) and cryoSPARC v3.2.0 (49). Drift-corrected averages of each movie were created using RELION’s implementation of MotionCor2 (50), and CTF parameters of each determined using Gctf (53). Both motion correction and CTF estimation were carried out on-the-fly (47). Initially, 1,368 particles were manually picked and used to train crYOLO v1.6.1 (52). This trained model was used for picking on all 227 movies, resulting in 117,630 particles. Particles were imported in RELION, extracted using a box size of 260 pixels and subsequently subjected to iterative rounds of reference-free 2D classification in cryoSPARC (mask diameter = 220 Å). After visual inspection, 88,412 particles were retained and used to generate four initial 3D volumes. Two of these models, which were characterized by continuous density and contained 20,487 and 21,879 particles respectively, were subsequently subjected to consensus 3D refinement with C1 symmetry, generating maps with reported global resolutions of 7.44 Å and 5.56 Å. Final resolutions were determined using the gold-standard Fourier shell correlation criterion (FSC = 0.143). The local resolution of the reconstructed maps was determined using the local resolution feature in cryoSPARC.

### Model building and refinement

An initial model for the cat BARD1^ARD-BRCTs^:H2AKc15ub nucleosomes complex was generated using ModelAngelo (54). Briefly, the protein sequences of cat BARD1^ARD-BRCTs^ (UniProt: C9IYG1), human histones (UniProt: P0C0S8, P33778, P68431 and P62805 for H2A.1, H2B, H3.1 and H4 respectively) and ubiquitin (Uniprot: P0CG48; aa 1-76) were used as inputs together with the final refined map (3.40 Å). To build an initial model of the Widom 601 DNA sequence, the cryo-EM structure of human nucleosomes (PDBid: 7XD1) (55) was rigid-body docked against the reconstructed density in ChimeraX v1.4 (56,57). The resulting model was then manually inspected and rebuilt using Coot v0.9.8.1 (58), and iterative rounds of real-space refinement were performed in Coot v0.9.8.1 and PHENIX v1.17.1 (59) using default parameters and secondary structure restraints. Isopeptide bond restraints were applied between Gly76 of ubiquitin and Lys15 of H2A. Amino acid residues that lacked unambiguous density were deleted or modelled up to their Cβ position while preserving sequence information. Gaps numbered arbitrarily were left where direct connectivity between secondary structure elements could not be determined. The overall quality of the model was assessed using MolProbity (60,61).

### AFM

All High-Speed AFM (HS-AFM) measurements were performed using a NanoRacer High-Speed AFM (Bruker) in amplitude modulation mode HS-AFM. All HS-AFM measurements were obtained in liquid and ambient temperature in an acoustic isolation housing on an active antivibration table using short cantilevers (USC-F1.2-k0.15, NanoWorld, Switzerland) with spring constant of 0.15 N m^−1^, resonance frequency of ∼0.6 MHz and a quality factor of ∼2 in buffer [25 mM HEPES pH 7.5, 75 mM NaCl, 5% (v/v) glycerol and 1 mM TCEP]. Samples were prepared using freshly cleaved mica substrates treated with Ni^2+^ to render the mica surface positively charged by incubating the mica with 100 mM NiCl_2_ for 2 min before rinsing with ultra-pure water. Di-nucleosomes were incubated with Ni^2+^ treated mica at 0.25 µg/ml for 3 min before rinsing with buffer [25 mM HEPES pH 7.5, 75 mM NaCl, 5% (v/v) glycerol and 1 mM TCEP] to remove unbound nucleosomes. No 6xHis tag were present for HS-AFM imaging, suggetsing attachment was driven by electrostatics only. For HS-AFM imaging of di- nucleosomes in the absence or presence of Human dStrepII-muGFP-BRCA1^ΔExon11^/6xHis-FL BARD1, BRCA1^Δ11^-BARD1 was added to and kept in the imaging solution at a concentration of 2.25 µg/ml. All images were obtained in imaging buffer containing 25 mM HEPES pH 7.5, 75 mM NaCl, 5mM NiCl_2_, 5% (v/v) glycerol and 1 mM TCEP. Localization AFM (LAFM) images of Di-nucleosomes were generated using 273 HS-AFM images of a single Di-nucleosome captured at 3 pixel/nm and processed with bicubic subpixel localization. HS-AFM movies and LAFM images were processed and analysed using custom written software in MATLAB (Matlab, Mathworks, Natick, MA, USA). Simulated AFM topography images were produced using an idealized AFM tip contacting 3D atomic coordinates within .pdb files for di- nucleosomes and lower resolution surfaces stored within .mrc files for BRCA1^Δ11^-:FL BARD1. Simulated topographies were generated using Mat-SimAFM software available at: https://github.com/George-R-Heath/Mat-SimAFM. Example images of BRCA1^Δ11^:FL BARD1 bridging across adjacent nucleosomes are shown in Supplementary Movie S1.

### Structure visualization

All structural models and surface representations depicted in Figures 4B, S4A, S5, & S7, were created using UCSF Chimera X v1.4 (56,57). The 3D FSC plot shown in Figure S5D was generated using the remote 3DFSC processing server, available at https://3dfsc.salk.edu/ (62). Interface measurements were perfomed using PISA (63) from BRCA1^RING^-BARD1^RING^ with nucleosome (21) and BARD1^ARD-BRCTs^ with nucleosome (22) structures.

## Results

### The BARD1 ARD and BRCT modules are specific readers of DNA damage-induced H2A ubiquitylation

To determine the ubiquitin-site specificity of the ankyrin repeat domain (ARD) and tandem BRCA1 C-terminal repeats (BRCTs) of BARD1, we expressed GST- and MBP-tagged BARD1^425-777^ (hereafter referred to as BARD1^ARD-BRCTs^) (Figure 1A, *Left*, Supplementary Figure S1A), and made nucleosomes harbouring ubiquitin marks at multiple biologically relevant sites (Figure 1A, *Right*, Figure 1B, Supplementary Figures S1B and C). We optimised a chemical alkylation approach used previously (39,44) to crosslink ubiquitin to specific histone residues using cysteine chemistry mimicking an isopeptide bond (see Material Methods, Supplementary Figure S1D). These included the DNA damage-associated marks H2AK13ub and H2AK15ub (31,64–66), the facultative heterochromatin mark H2AK119ub (67–69), BRCA1-BARD1 catalysed mark H2AK127ub (18,22,70) and the transcription elongation associated mark H2BK120ub (71,72). These are spatially distributed across the nucleosome, with H2AK13/15ub and H2BK120ub in close proximity on the back side of the nucleosome, and H2AK119ub and H2AK127ub on the opposite side adjacent to the DNA entry/exit sites (Figure 1B). GST-tagged BARD1^ARD-BRCTs^ protein immobilised on glutathione beads was able to enrich both H2AKc13ub and H2AKc15ub modified nucleosomes. By contrast, no interaction was observed for the nucleosomes bearing other ubiquitin marks (Figure 1B). Interestingly, despite H2BK120ub occupying a similar position on the nucleosome surface as H2AK13/15ub (∼9 Å separation), BARD1^ARD-BRCTs^ specifically reads the DNA damage-associated marks (Figure 1B), highlighting the exquisite BARD1 specificity to read DNA damage-induced ubiquitylation as part of the RNF168 signalling cascade. Intriguingly, ubiquitin specificity is highly salt dependant (Supplementary Figure S1E) with reduced complex formation for unmodified nucleosomes (upper band) at higher salt that is not affected in H2AKc15ub nucleosomes.

**Figure 1:**
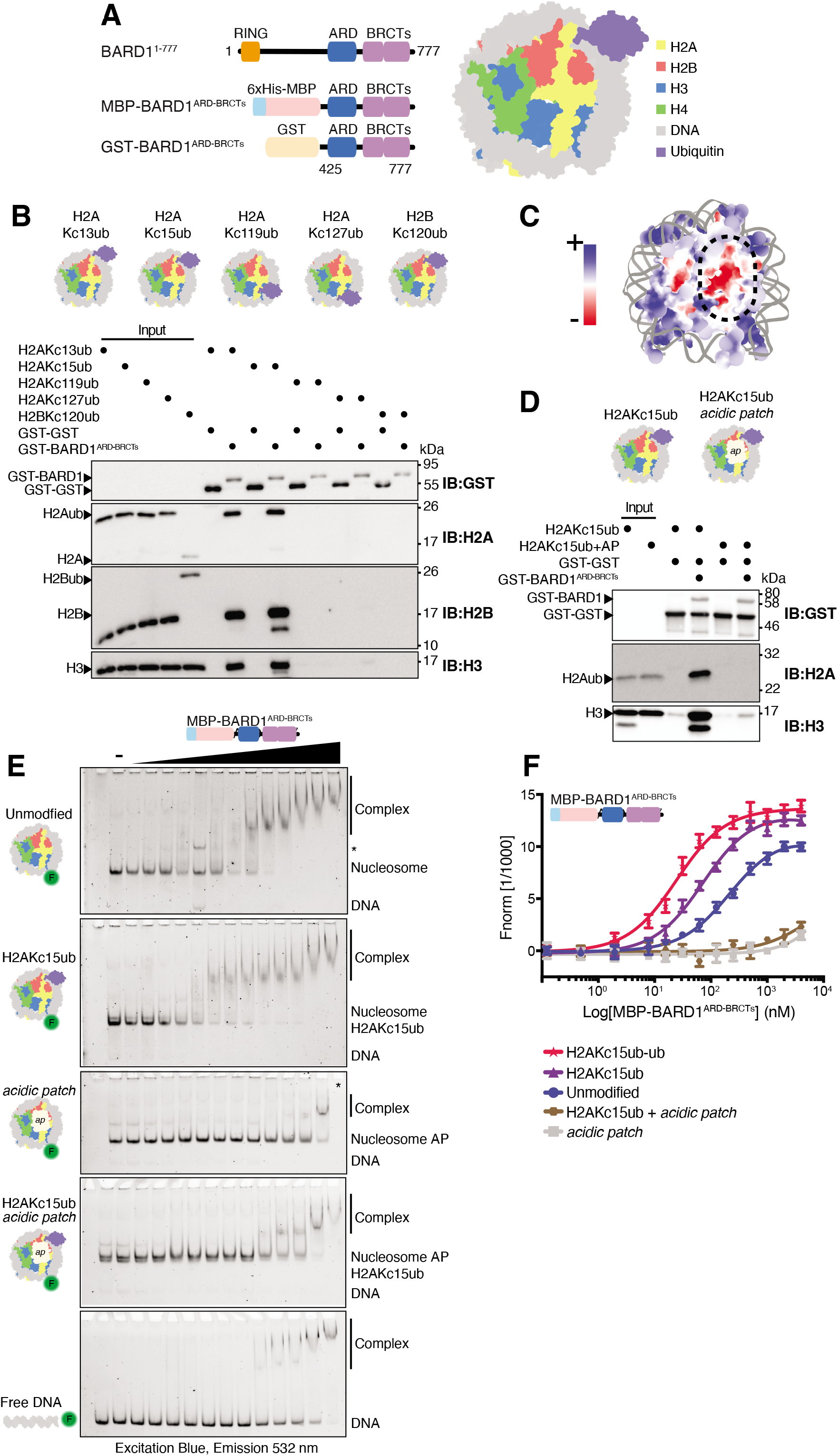
BARD1^ARD-BRCTs^ specifically recognises nucleosomes ubiquitylated at H2A Lys13/15, and is dependent on the nucleosome acidic patch. **A.** Schematics of the BARD1^ARD-BRCTs^ constructs (*Left*), and representation of a nucleosome ubiquitylated at H2A Lys 15 (*Right*). BARD1 = BRCA1-associated domain 1, RING = Really Interesting New Gene, ARD = Ankyrin Repeat Domain, BRCTs = BRCA1 C-terminus, MBP = Maltose Binding Protein tag, GST = Glutathione-S Transferase tag. The nucleosome model was made in USCF ChimeraX using nucleosome core particle structure (PDB ID: 1KX5) and ubiquitin (PDB ID: 1UBQ). **B.** Immunoblots from pull-down assay using GST-BARD1^ARD-BRCTs^ immobilised on glutathione affinity beads and incubated with the indicated purified recombinant ubiquitylated nucleosomes. Nucleosomes were chemically ubiquitylated at positions 13, 15, 119 & 127 in H2A, and 120 in H2B (representations shown above). **C.** Model of an unmodified nucleosome, coloured according to surface charge. Orientation of nucleosome is as in **A**. Blue indicates positive charge, red negative charge. The boxed region highlights the acidic patch sitting at the interface of H2A and H2B dimers. Model made in USCF ChimeraX using nucleosome core particle structure (PDB ID: 1KX5). **D.** Immunoblots from pull-down assay comparing the interaction of immobilised GST- BARD1^ARD-BRCTs^ with H2AKc15ub or H2AKc15ub/acidic patch (AP) mutant nucleosomes. For the acidic patch charge neutralising mutant, point mutations E61A, E91A, E92A were introduced in H2A and E105A in H2B. Both unmodified and mutant H2A were chemically ubiquitylated at position 15. For GST-BARD1 pull down assays equal amounts of GST-GST was also immobilised on the beads (lower band). A small degree of H3 degradation, overamplified by artifactual antibody recognition is highlighted with an asterisk. **E.** Electrophoretic mobility shift assays (EMSAs) comparing interaction of MBP-BARD1^ARD-BRCTs^ with nucleosome variants and free DNA. Nucleosomes were wrapped with 5’ FAM-labelled DNA (FAM = fluorescein) purified and limiting amounts (2.3 nM) were incubated with increasing concentrations (8-1920 nM) of MBP-BARD1^ARD-BRCTs^ protein. Lane marked with ‘–‘ indicates nucleosome alone. Complexes were resolved by native-PAGE and imaged using fluorescein filters. **F.** Amplitude-normalised microscale thermophoresis (MST) data assessing the binding affinity of 6xHis-MBP-BARD1^ARD-BRCTs^ for different nucleosome variants. Traces correspond to the titration of 6xHis-MBP-BARD1^ARD-BRCTs^ (0.122 nM-4000 nM) against 5’ FAM-labelled nucleosomes (at 20 nM each). Data are shown as average of three independent experiments +/-SEM. *K*_d_ values are summarised in Supplementary Table S1.

The H2A/H2B acidic patch is a negatively charged pocket on the face of the nucleosome (Figure 1C), and has been shown to be a crucial landing platform for many chromatin interacting proteins (73,74). To determine if BARD1^ARD-BRCTs^ interaction with nucleosomes is dependent on the acidic patch we used our chemical approach to generate nucleosomes carrying H2AKc15ub alone or in combination with charged neutralising acidic patch mutations on H2A and H2B (H2A^/E61A/E91A/E92A^ and H2B^E105A^) (Figure1D, Supplementary Figures S1F and G). BARD1^ARD-BRCTs^ showed greatly reduced binding to H2AKc15ub/acidic patch mutant nucleosomes using pull-down assays, electrophoretic mobility shift assays (EMSAs) and microscale thermophoresis (MST) (Figures 1D-F, Supplementary Figure S1H, Supplementary Table S1). Mutation of the acidic patch, either alone or in combination with H2AKc15ub, had a greater effect on binding compared to unmodified and H2AKc15ub nucleosomes, and reduced BARD1^ARD-BRCTs^ interactions to levels observed for DNA alone (Figures 1D-F, Supplementary Figure S1H, Supplementary Table S1). Taken together, these results demonstrate that BARD1^ARD-BRCTs^ acts as a specific reader of H2AK15ub nucleosomes and its nucleosome-recognition ability is largely dependent on an intact acidic patch, as also highlighted by recent structural studies (22,34).

### BRCA1-BARD1 complexes show increased affinity for nucleosomes

While the minimal fragment of BARD1^ARD-BRCTs^ is sufficient for interaction with modified mono-nucleosomes *in vitro*, the BRCA1-BARD1 complex includes other reported DNA and chromatin interacting regions (22,35,75). To determine if BRCA1-BARD1 has higher affinity for nucleosomes compared to BARD1^ARD-BRCTs^ alone, we co-purified a complex comprising the common BRCA1 ΔExon11 splice variant and full-length BARD1 (Flag-BRCA1^Δ224-1365^:6xHis- BARD1^1-777^, hereafter termed BRCA1^Δ11^:BARD1) (Figure 2A, Supplementary Figure S2A). We found this was the largest fragment of BRCA1-BARD1 complex that was stably expressed and purified at sufficient levels for our assays. In MST assays, BRCA1^Δ11^:BARD1 bound unmodified nucleosomes with twice the affinity compared to the isolated BARD1^ARD-BRCTs^ protein (Figure 1F, Figure 2B, Supplementary Table S1). Inclusion of the H2AKc15ub mark again doubled the affinity as compared to BARD1^ARD-BRCTs^, and mutation of the nucleosome acidic patch significantly reduced binding to both unmodified and H2AKc15ub nucleosomes (Figure 1F, Figure 2B, Supplementary Table S1). These data demonstrate that the reconstituted BRCA1^Δ11^:BARD1 complex has enhanced affinity for nucleosome substrates compared to BARD1^ARD-BRCTs^, confirming that additional regions in BRCA1 and/or BARD1 mediate interactions with the histone octamer or nucleosomal DNA.

**Figure 2:**
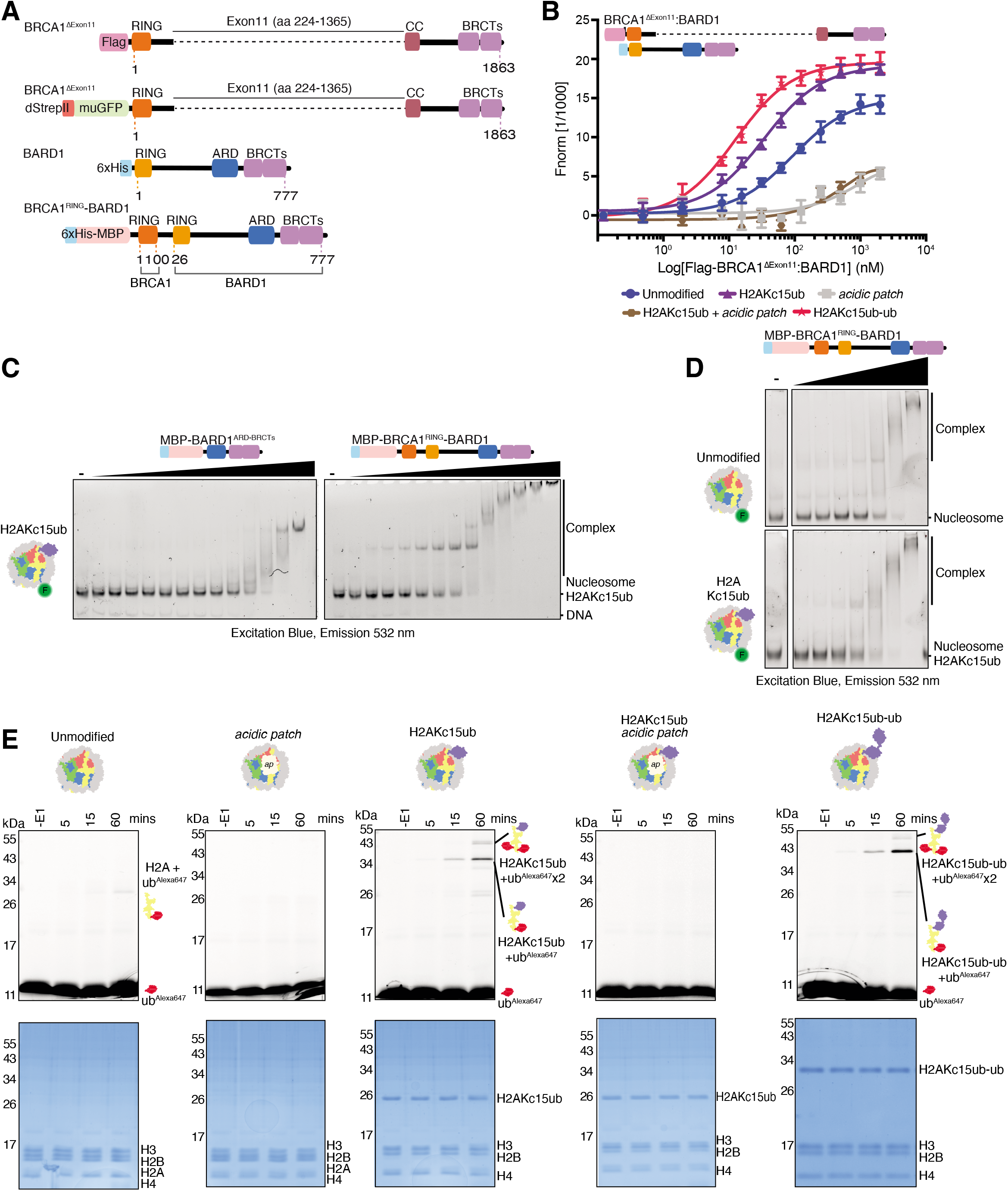
BRCA1-BARD1 complexes show increased affinity for nucleosomes and have E3 ligase activity. **A.** Schematics of the BRCA1^Δ11^:BARD1 co-expression construct and the BRCA1^RING^-BARD1 genetically fused construct. BRCA1 = Breast cancer type 1 susceptibility protein, BARD1 = BRCA1-associated domain 1, RING = Really Interesting New Gene, CC = coiled-coil, BRCTs = BRCA1 C-terminus, ARD = Ankyrin Repeat Domain, dStrepII = Double StrepII tag, muGFP = monomeric ultra-stable GFP tag, MBP = Maltose Binding Protein tag. **B.** Amplitude-normalised MST data assessing the binding affinity of Flag-BRCA1^Δ11^:6xHis- BARD1 for indicated nucleosome variants. Traces correspond to the titration of Flag- BRCA1^Δ11^:6xHis-BARD1 (0.122 nM-2000 nM) against 5’ FAM-labelled nucleosomes (at 20 nM each). Data are shown as average of three independent experiments+/-SEM.. *K*_d_ values are summarised in Supplementary Table S1. **C.** EMSA experiments comparing the relative affinity of 6xHis-MBP-BARD1^ARD-BRCTs^ and the larger 6xHis-MBP-BRCA1^RING^-BARD1 to H2AKc15ub modified nucleosomes with 5’ FAM-labelled DNA. Protein concentration range was 1-2100 nM. **D.** EMSA experiments testing BRCA1^RING^-BARD1 binding to unmodified and H2AKc15ub mono-nucleosomes. Nucleosomes were wrapped with 5’ FAM-labelled DNA, and increasing concentrations (8-512 nM) of fused BRCA1^RING^-BARD1 complex were used. A higher salt concentration than **C** (150 mM as opposed to 75 mM in **C**) was used. Complexes were resolved by native-PAGE and imaged for fluorescein. **E.** Ubiquitylation assays using BRCA1^RING^-BARD1 and recombinant nucleosomes, assessing the role of the acidic patch (AP, H2A^E61A/E91A/E92A^ and H2B^E105A^) mutant and H2A K15 ubiquitylation on E3 ligase activity. Samples were taken prior to addition of E1 (-E1) and at 5, 15 and 60 minutes time points, and quenched by addition of 2x SDS loading buffer. Samples were resolved on SDS-PAGE gels and directly imaged measuring Alexa647 signal, before staining with Coomassie. Top panel shows the Alexa647 signal (Excitation Red light, emission 700 nm/50 nm), and lower panel a Coomassie stain of the total protein.

The BRCA1-BARD1 RING domains and the BARD1 ARD-BRCT modules have been implicated in BRCA1-BARD1 recruitment as well as E3 ligase enzymatic activity (36,76) (11,18,21,22,32–34,77,78). This wealth of information highlights the central role of BARD1 in supporting BRCA1-BARD1 activity and chromatin recruitment. To understand how these multiple BRCA1-BARD1 modules function using a minimally reconstituted system, we purified a genetically fused protein containing near full-length BARD1 with the RING domain from BRCA1 (6xHis-MBP-BRCA1^1-100^-BARD1^26-777^, hereafter termed BRCA1^RING^-BARD1) (Figure 2A, Supplementary Figure S2A). This fragment purified more readily and was more stable than larger assemblies while behaving biochemically similarly. EMSA assays showed a higher affinity of BRCA1^RING^-BARD1 for H2AKc15ub nucleosomes compared to BARD1^ARD-BRCTs^ alone (Figure 2C), and the fused BRCA1^RING^-BARD1 protein bound with similar affinity as the larger BRCA1^Δ11^:BARD1 (Supplementary Figure S2B). Consistent with the results obtained with BRCA1^Δ11^:BARD1 (Figure 2B, Supplementary Table S1), BRCA1^RING^-BARD1 also exhibited stronger affinity for H2AKc15ub modified versus unmodified nucleosomes (Figure 2D), and mutation of the acidic patch diminished BRCA1^RING^-BARD1 interactions with H2AKc15ub nucleosomes (Supplementary Figure S2C). Collectively, these data show that larger BRCA1-BARD1 complexes have increased binding affinity to nucleosomes than BARD1^ARD-BRCTs^ alone. This higher affinity interaction is a result of nucleosome binding by the BRCA1-BARD1 RINGs, the BARD1 ARD-BRCTs and possibly the unstructured region between the BARD1 RING and ARD domains.

### BRCA1-BARD1 E3 ligase activity is dependent on the nucleosome acidic patch

Previous studies have shown the BRCA1^RING^-BARD1^RING^ heterodimer is an active E3 ligase targeting H2A C-terminal lysine residues 125/127/129 (18,20–22). In line with these findings, we were able to recapitulate ubiquitylation H2A on unmodified nucleosomes using BRCA1^RING^-BARD1 (Figure 2E, Supplementary Figure S2D), observing shifts in electrophoretic mobility of fluorescent ubiquitin in SDS-PAGE gels corresponding to H2A becoming modified by ubiquitin. Bands running at ∼26 kDa correposnd to single H2A C-terminal ubiquitylation in unmodified nucleomes. For the already larger H2AKc15ub, bands above ∼34kDa and ∼43kDa correposnd to a single or second ubiquitin added at C-terminal lysines. Interestingly, activity was higher on H2AKc15ub modified nucleosomes, suggesting that increased recruitment driven by interactions between the BARD1^ARD-BRCTs^ and ubiquitylated H2A K15 stimulates productive ubiquitylation by the BRCA1-BARD1 RINGs on the H2A C-terminal tail. Indeed, the efficiency in E3 ligase activity can be used as a proxy for BRCA1-BARD1 heterodimer binding to its nucleosomal substrates. Consistent with binding assays, mutation of the acidic patch on unmodified or H2AKc15ub nucleosomes abrogated H2A ubiquitylation activity (Figure 2E). Overall, BRCA1^RING^-BARD1 is an active E3 ligase targeting H2A, stimulated by H2A K15ub and dependent on an intact nucleosome acidic patch.

### Lysine 63-linked ubiquitin chains do not preclude BRCA1-BARD1 binding to nucleosomes

E3 ligases RNF8 and RNF168 promote the synthesis of lysine 63 (K63)-linked ubiquitin chains neighbouring to DNA DSBs (31,79), and previous studies have suggested that H2A may be a target for these chains (65,80). It was recently shown that BARD1 binding prevents extension of K63-linked ubiquitin chains on H2AK15ub nucleosomes, with Lys 63 on the acceptor ubiquitin in close proximity to the BARD1 BRCTs (Figure 3A) (22). However, how ubiquitin chains conjugated at H2AK15ub affects the recruitment of BARD1 to DSBs, and the effect they may have on BRCA1^RING^-BARD1^RING^ heterodimer activity, is not known.

**Figure 3:**
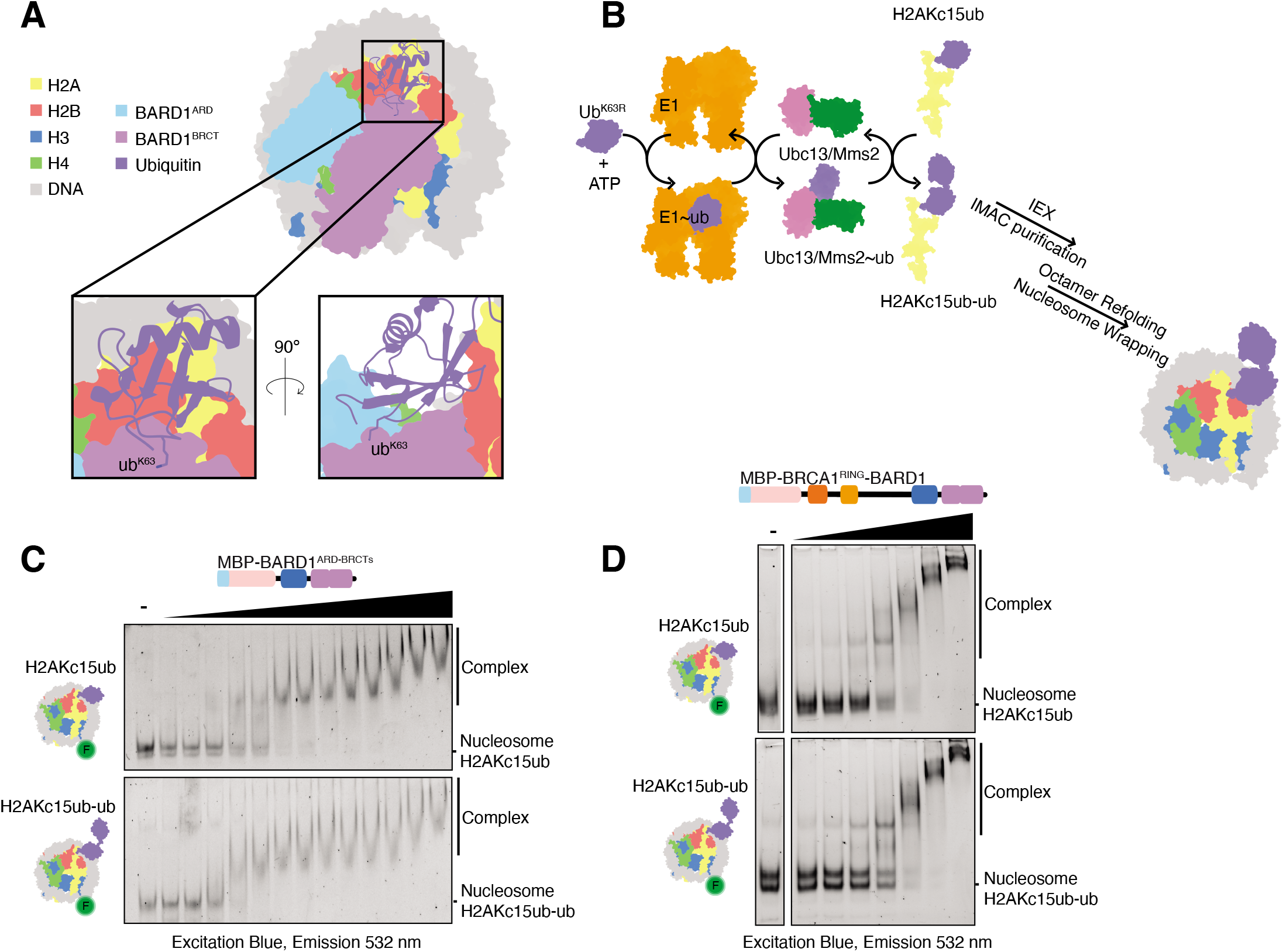
K63-linked ubiquitin chains do not preclude BARD1^ARD-BRCTs^ binding to nucleosomes. **A.** Structure of the BARD1 ARD-BRCT regions bound to H2A K15 ubiquitylated nucleosomes (PDB ID: 7LYC), with magnified view of K63 residue on ubiquitin. ARD = Ankyrin Repeat Domain, BRCT = BRCA1 C-terminus, Ubiquitin = Ub. **B.** Schematic of the ubiquitylation assays used to make H2AKc15ub-ub^K63R^ histones. Chemically ubiquitylated H2AKc15ub histone was combined in an assay buffer with ATP, 6xHis-TEV-Ubiquitin^K63R^ and E2 enzymes Ubc13/Mms2. Reactions were initiated by addition of E1 enzyme (*Left*). Schematic representation of nucleosome with K63-linked di-ubiquitin at H2A position K15 (*Right*). **C-D.** EMSA experiments comparing interactions between H2AKc15ub and H2AKc15ub-ub^K63R^ nucleosomes wrapped with 5’ FAM-labelled DNA and increasing concentrations of BARD1^ARD-BRCTs^ (8-1920 nM) (**C**) or BRCA1^RING^-BARD1 (8-512 nM) (**D**) proteins. Complexes were resolved by native-PAGE and imaged for fluorescein.

To determine the effect K63-linked ubiquitin chains have on BRCA1-BARD1 nucleosome binding and E3 ligase activity, we generated K63-linked di-ubiquitin chains tethered to H2A at residue 15 by combining chemical and enzymatic ubiquitylation (H2AKc15ub-ub). The chemically crosslinked H2AKc15ub protein as a substrate and the K63- specific E2 pair Ubc13/Mms2 (81–83) was used to add a chain capping Ub^K63R^ mutant (84), thus generating a di-ubiquitylated H2AKc15ub-ub^K63R^ histone that could be purified and assembled into nucleosomes (Figure 3B, Supplementary Figure S3). Surprisingly, the minimal BARD1^ARD-BRCTs^ protein displayed a higher affinity for H2AKc15ub-ub^K63R^ over H2AKc15ub nucleosomes (Figure 1F, Figure 3C, Supplementary Table S1). Likewise, both BRCA1^Δ11^:BARD1 and BRCA1^RING^-BARD1 showed enhanced binding to H2AKc15ub-ub nucleosomes compared to mono-ubiquitylated nucleosomes (Figure 2B, Figure 3D, Supplementary Table S1).

To determine how K63-linked ubiquitin chains on H2A K15 affect BRCA1^RING^- BARD1^RING^ E3 ligase activity, we performed ubiquitylation assays on H2AKc15ub-ub nucleosomes. Consistent with binding experiments, BRCA1^RING^-BARD1 ubiquitylated H2AKc15ub-ub and H2AKc15ub nucleosomes at similar levels, and its E3 ligase activity was much higher on both of these substrates than on unmodified nucleosomes (Figure 2E, Supplementary Figure 3E). Overall, these results show that K63-linked ubiquitin chains on H2A K15 are not a barrier to BRCA1-BARD1 chromatin interaction, nor an inhibitor of their enzymatic activity.

### BRCA1-BARD1 does not readily contact both faces of a mono-nucleosome

Recent single particle cryogenic-electron microscopy (cryo-EM) structures of the BRCA1^RING^-BARD1^RING^ heterodimer and the BARD1^ARD-BRCTs^ have provided detailed insights on how these separate domains engage nucleosomes, thereby promoting BRCA1-BARD1 recruitment to its substrates (21,22,34). However, how these multiple nucleosome-binding entities concomitantly arrange on chromatin, and how this is regulated in the context of the full-length BRCA1-BARD1 complex, is still unclear. Sterically, the BRCA1^RING^-BARD1^RING^ and BARD1^ARD-BRCTs^ cannot simultaneously bind to the same face of a nucleosome, as both modules mediate contacts with residues localized in the H2A/H2B acidic patch (Supplementary Figure S4A). The nucleosome itself is pseudo-symmetrical, with two identical protein solvent exposed surfaces. One hypothesis for BRCA1-BARD1 engagement is that these two nucleosome-binding modules interact and ‘clamp’ concurrently on either face of a single nucleosome. In this model, the BRCA1^RING^-BARD1^RING^ heterodimer would bind to one side of the nucleosome particle while the BARD1^ARD-BRCTs^ interact with the opposite interface (Figure 4A) (85).

**Figure 4:**
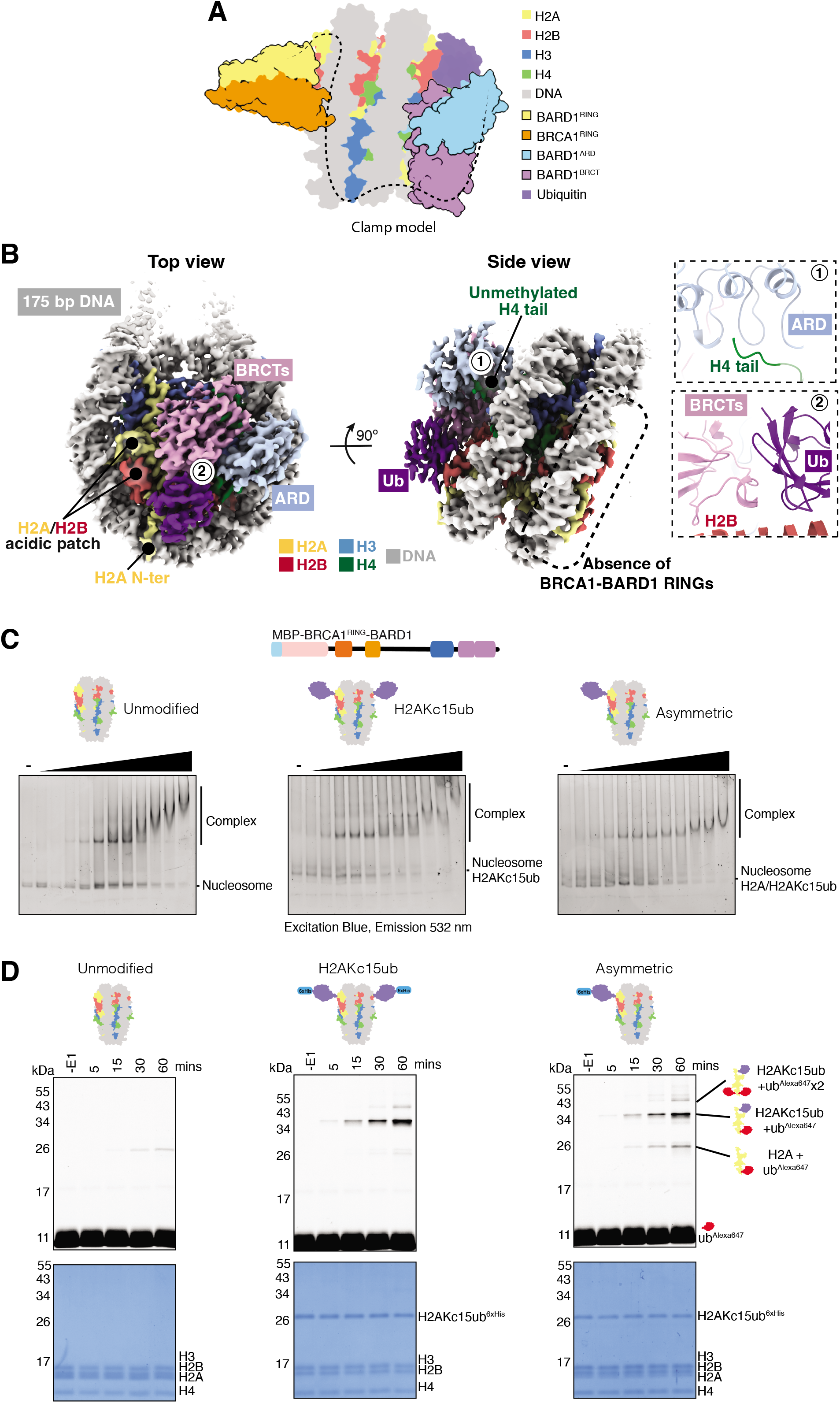
BRCA1-BARD1 does not readily ‘clamp’ over mono-nucleosomes. **A.** Schematic representation of the proposed ‘clamp’ model for BRCA1-BARD1 binding to mono-nucleosomes. In this model, the BARD1 ARD-BRCTs bind to H2AK15ub on one side of the nucleosome, guiding the BRCA1-BARD1 RING domains to the opposite surface. The central unstructured region in BARD1 may contact nucleosomal DNA. RING = Really Interesting New Gene, ARD = Ankyrin Repeat Domain, BRCTs = BRCA1 C-terminus. **B.** 3.40 Å cryo-EM map (at a contour level of 0.005) of BARD1^ARD-BRCTs^ bound to H2AKc15ub nucleosomes, shown in two orientations and coloured by corresponding chain. Absence of density for the BRCA1-BARD1 RING domains on the opposite nucleosome interface is indicated with a dashed black rectangle. Insets show interactions between the BARD1 ARD and the unmethylated H4 tail, and between the BARD1 BRCTs and H2AK15-linked ubiquitin. **C.** EMSA experiments comparing binding of MBP-BRCA1^RING^-BARD1 to unmodified (*Left*), symmetric H2AKc15ub (*Middle*; two copies of ubiquitylated H2A) and asymmetric H2AKc15ub/H2A unmodified (*Right*; one copy of ubiquitylated H2A) nucleosomes. Nucleosomes were wrapped with 5’ FAM-labelled DNA, and increasing concentrations (10- 200 nM) of MBP-BRCA1^RING^-BARD1 protein were incubated. Complexes were resolved by native-PAGE and imaged for fluorescein. **D.** Ubiquitylation assays using BRCA1^RING^-BARD1 and recombinant nucleosomes. Nucleosomes were either unmodified (*Left*), symmetrically ubiquitylated (*Middle*) or asymmetrically ubiquitylated on only one H2A protomer per octamer (*Right*). Samples were taken prior to addition of E1 (-E1) and at 5, 15, 30 and 60 minute time points, and quenched by addition of 2x SDS loading buffer. Samples were resolved on SDS-PAGE and imaged for Alexa647 signal before staining with Coomassie. Top panel shows the Alexa647 signal, and lower panel a Coomassie stain of the total protein. Higher migrating bands correspond to ubiquitylated species, with the highest band being multi-mono ubiquitylated.

To address how a fully assembled larger fragment of BRCA1-BARD1 complex interacts with its substrates, we determined the structure of BRCA1^Δ11^BARD1 bound to H2AKc15ub nucleosomes. We obtained a map at a global resolution of 3.4 Å with clear density for the coin-shaped nucleosome core particle, thus allowing *de novo* model building of the histone octamer and unambiguous fitting of core nucleosome DNA sequence (40) (Figure 4B, Supplementary Figure S5A-D, Supplementary Table S2). There was instead poorer density for the 15 bp DNA linker arms (Supplementary Figure S5B), suggesting these are flexible and not tethered or stabilised by the BRCA1^Δ11^-BARD1 complex. Interestingly, on one face of the nucleosome we observed well-ordered density for the covalently H2A-linked ubiquitin and extra density attributable to the BARD1 ARD-BRCTs (Figure 4B). This minimal nucleosome-interacting module was arranged in a similar manner as the analogous BARD1^ARD-BRCTs^- nucleosome structures recently determined after glutataldehyde chemical crosslinking (Supplementary Figure S5E) (22,34) recently determined after glutaraldehyde chemical crosslinking. In agreement with these models and our biochemistry, we observed direct interactions between the BARD1 ARD and the unmethylated H4 tail as well as the tandem BRCT domains contacting the nucleosome acidic patch and H2A-linked ubiquitin (Figure 4B). Surprisingly, we could not visualize significant additional density on the opposite face of the nucleosome, where we would have expected the BRCA1-BARD1 RING domains to bind in the ‘clamp’ model. At this interface, we could only identify diffuse density localised over the H2A N-terminal tail which is consistent with unbound H2AKc15ub, as observed previously (Supplementary Figure S5B) (39).

To investigate this unexpected result, we strove to test the clamping model biochemically by creating asymmetrically ubiquitylated nucleosomes. Recent studies demonstrated that the BARD1^ARD-BRCTs^ bind with higher affinity to H2AK15ub nucleosomes than the BRCA1^RING^-BARD1^RING^ heterodimer (21,22), suggesting the ARD and BRCT modules drive BRCA1-BARD1 complex recruitment to chromatin. We hypothesized that by producing asymmetric nucleosomes, containing one ubiquitylated (H2AK15ub) and one unmodified H2A on the other face of the nucleosome, we could provide a substrate where the BARD1^ARD-BRCTs^ would preferentially bind on the ubiquitylated H2A surface thus leaving the unmodified H2A interface available for binding by the BRCA1^RING^-BARD1^RING^ heterodimer. This substrate allows us to assess BRCA1-BARD1 binding and activity, as we would expect to observe higher E3 ligase activity on the unmodified H2A of the asymmetric nucleosome than on the H2AKc15ub side. We generated asymmetric nucleosomes using a tag and purification approach (Supplementary Figure S4B-D) (46), and we could distinguish which face of the nucleosome particle was ubiquitylated based on difference in SDS-PAGE mobility using labelled ubiquitin (H2AKc15ub+K125/127/129ub^Alexa647^ versus H2A+K125/127/129ub^Alexa647^).

By EMSA assays, BRCA1^RING^-BARD1 showed a marginally reduced binding to these asymmetric mono-nucleosomes compared to H2AKc15ub substrates (Figure 4C). In a ubiquitylation assay there was activity on both the modified and unmodified H2A (Figure 4D). However, contrary to our hypothesis the activity on the unmodified face of the nucleosome did not increase appreciably (Figure 4D), although higher than for fully unmodified nucleosome. These data suggest that while BRCA1-BARD1 can clamp and ubiquitylate the unmodified nucleosome surface this is not favoured over the ubiquitylated face. These results, combined with the cryo-EM structure, therefore suggest the ‘clamp’ model may not represent the preferred BRCA1-BARD1 nucleosome binding mode.

### BRCA1-BARD1 bridges between adjacent nucleosomes

As the ‘clamp’ hypothesis did not appear to be favoured by our biochemical and structural experiments, we next sought to explore a BRCA1-BARD1 ‘bridge’ model of nucleosome binding. In this scenario, the BARD1^ARD-BRCTs^ would bind to one nucleosome while the BRCA1^RING^-BARD1^RING^ heterodimer spans across to an adjacent nucleosome, effectively bridging between these nucleosomes (Figure 5A) (85). To test this hypothesis, we produced recombinant di-nucleosomes with a 30 bp linker DNA, and performed EMSA and ubiquitylation assays. In EMSA experiments using H2AKc15ub nucleosome variants, BRCA1^RING^-BARD1 had a higher binding affinity for di-nucleosomes than mono-nucleosomes (Figure 5B). As previously observed for mono-nucleosome substrates, binding to di-nucleosomes was also driven by interactions with an intact acidic patch and enhanced by ubiquitylation at H2A Lys- 15 (Figure 5B, Supplementary Figure S6A). Consistent with binding assays, we observed stronger ubiquitylation activity when using di-nucleosomes as a substrate compared to mono- nucleosomes, for both unmodified and H2AKc15ub, despite an equimolar amount of available substrate H2A (Figure 2E, Figure 5C). Likewise, BRCA1^RING^-BARD1 E3 ligase activity on di- nucleosomes was also dependent on an intact acidic patch (Supplementary Figure S6B). These data therefore suggest that di-nucleosomes represent a preferential substrate for BRCA1-BARD1 compared to mono-nucleosomes.

**Figure 5:**
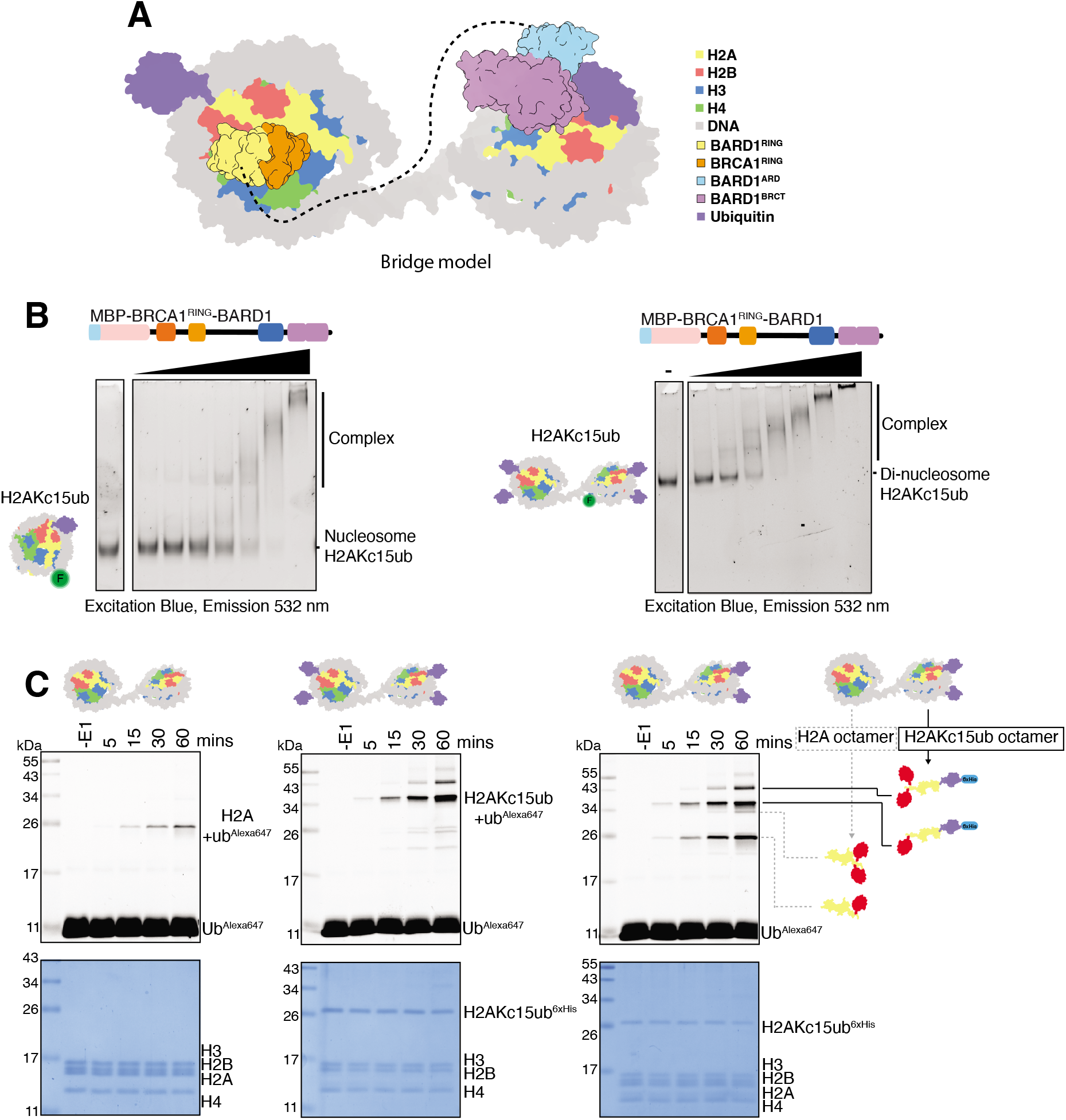
BRCA1-BARD1 can bridge between adjacent nucleosomes. **A.** Schematic representation of the proposed ‘bridge’ model for BRCA1-BARD1 binding to di- nucleosomes. In this model, the BARD1 ARD-BRCTs bind to H2AK15ub on one nucleosome surface, guiding the BRCA1-BARD1 RING domains to the interacting surface on an adjacent nucleosome particle. The central unstructured region of BARD1 could interact with linker DNA between the two nucleosomes to further strengthen the complex. RING = Really Interesting New Gene, ARD = Ankyrin Repeat Domain, BRCTs = BRCA1 C-terminus. **B.** EMSA experiments comparing binding of BRCA1^RING^-BARD1 to recombinant mono- and di-nucleosomes. Nucleosomes were wrapped with 5’ FAM-labelled DNA, and increasing concentrations (8-512 nM) of MBP-BRCA1^RING^-BARD1 protein were used. Complexes were resolved by native-PAGE and imaged for fluorescein. **C.** Ubiquitylation assays using BRCA1^RING^-BARD1 and recombinant di-nucleosomes with different H2A ubiquitylation status. Comparison between unmodified (*Left*), fully ubiquitylated (*Middle*) or singly ubiquitylated on only one nucleosome (*Right*). Samples were taken prior to addition of E1 (-E1) and at 5, 15, 30 and 60 minutes time points, and quenched by addition of 2x SDS loading buffer. Samples were resolved on SDS-PAGE and imaged for Alexa647 signal, before staining with Coomassie. Top panel shows the Alexa647 signal and lower panel a Coomassie stain of the total protein.

To test the ‘bridge’ hypothesis, we produced heterotypic di-nucleosomes, again levying the separation of ubiquitin binding preference between the BRCA1-BARD1 RINGs and BARD1 ARD-BRCTs. We created heterotypic nucleosomes whereby one nucleosome particle has both H2AK15 ubiquitylated and the other is unmodified. Using the same strategy and methodology already employed for the reconstitution of heterotypic mono-nucleosomes (46) (Supplementary Figure S6C) we were able to separate different H2AKc15 ubiquitylation states of the di-nucleosome substrate based on differential elution from affinity chromatography resins (Supplementary Figures S6D and E). If BRCA1-BARD1 could bridge across di- nucleosomes, the the ARD-BRCT module would preferentially bind the H2AK15ub protomer.

As such we anticipate increased catalytic activity on the normally poorly ubiquitylated unmodified H2A. Indeed, we observed an increase in activity on unmodified H2A in the heterotypic di-nucleosomes compared to the fully unmodified symmetric nucleosomes (Figure 5C, Supplementary Figure S6F), in contrast to the weak activity seen with asymmetric H2AK1c15ub/H2A mono-nucleosomes (Supplementary Figure S4D). This suggests that interactions between the BARD1^ARD-BRCTs^ with ubiquitylated H2A K15 on one nucleosome surface facilitate the binding of the BRCA1^RING^-BARD1^RING^ heterodimer to the corresponding interface of the neighbouring nucleosome particle.

### BRCA1-BARD1 samples multiple conformations

To gain further insights into our ‘bridge’ model, we initially sought to investigate the structural features of BRCA1-BARD1 and the interactions occurring within this complex. We obtained two low-resolution cryo-EM maps of BRCA1-BARD1 in different conformational states that highlighted a high degree of structural flexibility (Figure 6A, Supplementary Figure S7, Supplementary Table S2). In line with a previous report (86), the “open state” BRCA1- BARD1 map adopts an extended crescent like shape with two lobes at either side of the connected density (Figure 6A, *Left*). By contrast, the “closed state” map suggests potential interactions between separate domains or regions within BRCA1-BARD1 (Figure 6A, *Right*). Real-time single molecule observations using high-speed atomic force microscopy (HS-AFM) also showed a high degree of BRCA1-BARD1 flexibility and its remarkable ability to attain different conformations (Figure 6B). These analyses suggest high conformational dynamics within BRCA1-BARD1, with the complex adopting different states that may be important for enzyme activity and/or substrate recruitment.

**Figure 6:**
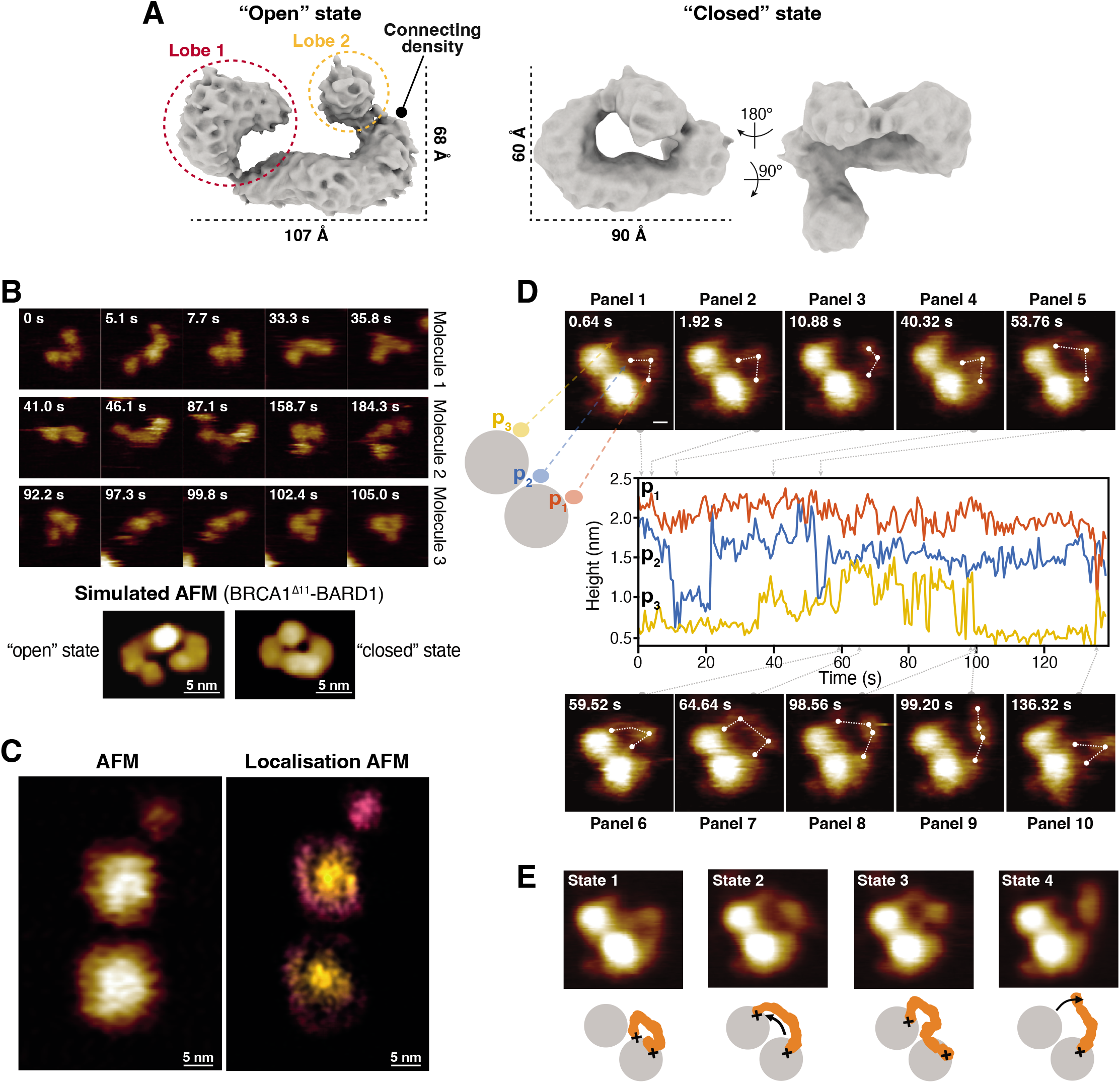
BRCA1-BARD1 is flexible and can be visualised interacting with di- nucleosomes using atomic force microscopy. **A.** Cryo-EM maps of BRCA1^Δ11^:BARD1 complex in the “open” (*Left*; contour level = 0.317) and “closed” (*Right*; contour level = 0.307) states, with map dimensions indicated. **B.** High speed-atomic force microscopy (HS-AFM) of BRCA1^Δ11^:BARD1, showing three representative BRCA1^Δ11^:BARD1 molecules captured over time. All images were taken from a single movie (*Top*). AFM simulation of BRCA1^Δ11^:BARD1 “open” (*Left*) and “closed” (*Right*) states. Tip radius and sampling values used in simulation AFM were 1 nm and 3 pixels/nm, respectively (*Bottom*). **C.** AFM (*Left*) and localisation AFM (LAFM; *Right*) of unmodified di-nucleosome particles in the absence of BRCA1^Δ11^:BARD1. **D.** HS-AFM of BRCA1^Δ11^:BARD1 in the presence of unmodified di-nucleosomes, with height analysis at three different contact points (p1, p2, p3). The sampling value was 2.56 pixels/nm, and the speed was 1.5625 fps with bi-directional scanning. **E.** Averaged AFM images of four identified states from **D**, with proposed models of molecular arrangements. The number of frames averaged per state were six, eight, seven and 16 respectively.

### Real-time observation of BRCA1-BARD1 binding to nucleosomes

To validate our biochemical and structural analyses, and independently visualize BRCA1-BARD1 interactions with di-nucleosomes, we performed HS-AFM experiments using di-nucleosomes and the BRCA1^Δ11^:BARD1 complex. We initially recorded independent movies for di-nucleosomes, and we took advantage of a published nucleosome structure (PDBid: 7PF4) (87)) and of our low-resolution cryo-EM maps (Figure 6A) to generate AFM simulated models for data interpretation (Figure 6B, Supplementary Figure S8A). We could visualise static nucleosome particles in the absence of BRCA1-BARD1, and localization AFM (LAFM) image reconstruction (88) provided a clear visual of DNA and histone proteins (Figure 6C, Supplementary Figure S6B).

Encouraged by these data, we performed real-time measurements of di-nucleosomes after adding BRCA1^Δ11^:BARD1. We were able to visualize di-nucleosome particles with three additional main extra heights (p1-p3) that were not visible in the isolated di-nucleosomes dataset and that we attributed to BRCA1^Δ11^:BARD1 (Figure 6D, Supplementary Figures S6C and D). Two heights occurred on one nucleosome particle rapidly, and these were centred on p1 and p2 contact points (Figure 6D, *panel 1,* Supplementary Movie 1). While the residency height for p1 remained stable over the time course of the experiment, the heights at p2 and p3 appeared more dynamic and could lift (p2) and bridge across to the adjacent nucleosome particle (p3) in between 50-70 seconds (Figure 6D, *panels 2-7*). Averaging of selected frames revealed at least four potential BRCA1-BARD1 engaged states, highlighting the large degree of flexibility and multivalent interactions with di-nucleosomes (Figure 6E), with the first contact point (p1) having the highest residency time. These data are consistent with a model for BRCA1-BARD1 recruitment to di-nucleosomes characterized by four possible states: association with one nucleosome particle (Figure 6E, *state 1*) and ‘bridge’ movement across the adjacent nucleosome by contacting the linker DNA (Figure 6E, *states 2 and 3*), prior to final disengagement from this second nucleosome moiety (Figure 6E, *state 4*).

Collectively, real-time observation of BRCA1-BARD1 engaging di-nucleosomes are consistent with our biochemical and enzymology data suggesting that BRCA1-BARD1 favours bridging of adjacent nucleosomes.

### BRCA1-BARD1 bridging navigates nascent chromatin by bridging and tolerating of H4K20-methylation

BRCA1-BARD1 is recruited to damage-adjacent post-replicative chromatin, and the timing of this recruitment is coordinated to promote HDR in the S/G2 phase of the cell cycle when a sister chromatid is present. Mechanistically, this is in part mediated via chromatin marks and requires BARD1^ARD-BRCTs^ recognition of both H2AK15ub (33) as well as of nascent chromatin which lack H4K20 methylation (H4K20me0) (32) (Figure 7A). In contrast, H4K20 di-methylation (H4K20me2) blocks BARD1^ARD-BRCTs^ binding and is spread throughout the genome as a marker of chromatin age (89).

**Figure 7:**
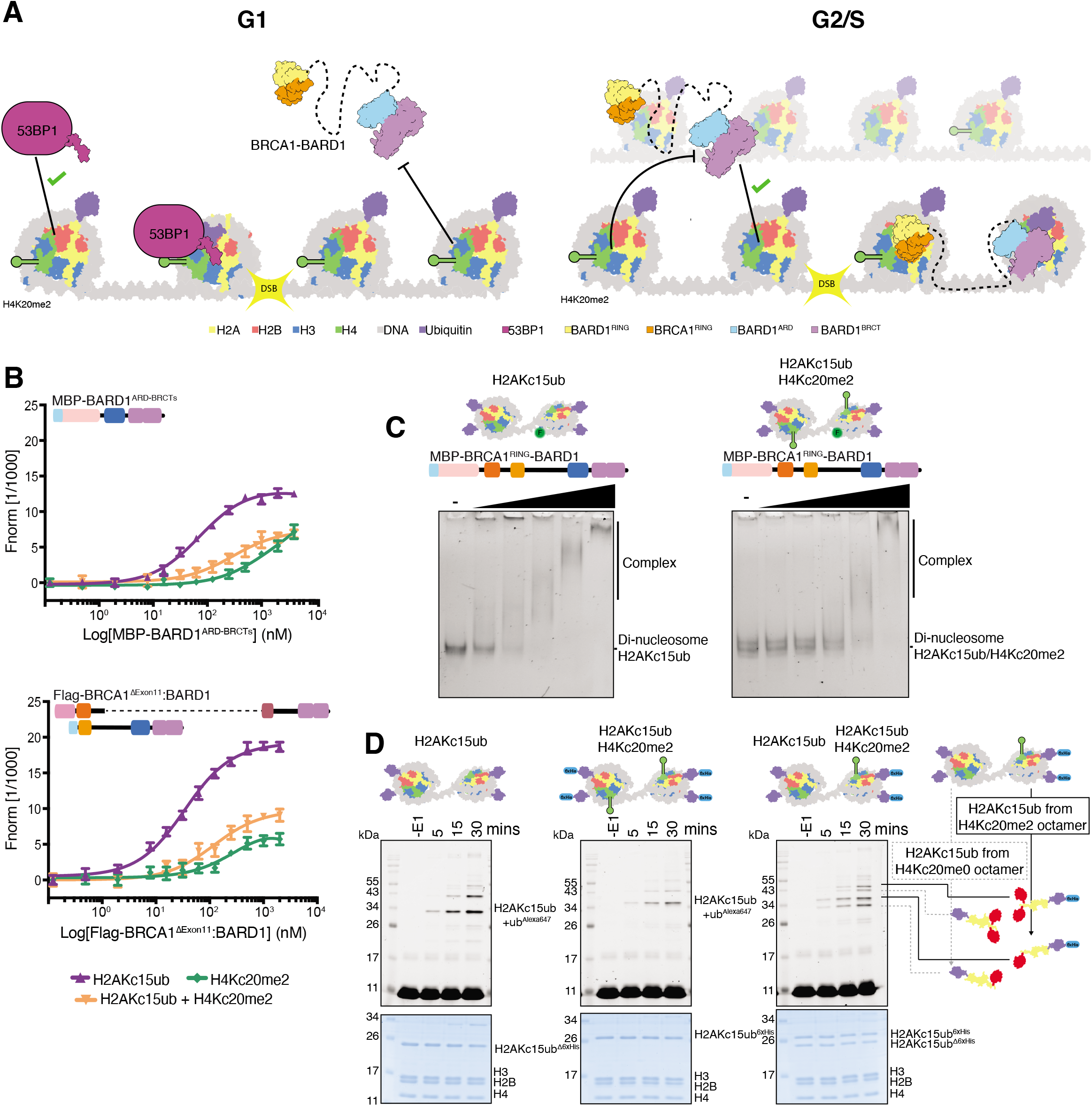
BRCA1-BARD1 nucleosome bridging enables H2A C-terminal ubiquitylation of H4K20me2-containing nucleosomes. **A.** Proposed model of BRCA1-BARD1 chromatin recruitment at DNA damage sites. The majority of H4 tails are di-methylated at position 20 (H4K20me2) in G1 phase of cell cycle. This methylation precludes BARD1 binding and promotes 53BP1 binding. After replication of sister chromatids in S and G2 phases, newly deposited H4 is unmethylated at H4K20, essentially leaving every other nucleosome retaining the methylation. This unmethylated H4 allows BARD1 to bind and recruits BRCA1-BARD1 RING domains to all nucleosomes around the break site, despite the presence of H4K20me2 marks on half of the nucleosomes. RING = Really Interesting New Gene, ARD = Ankyrin Repeat Domain, BRCTs = BRCA1 C-terminus. **B.** Amplitude-normalised MST data assessing the binding affinities of MBP-BARD1^ARD-BRCTs^ (*Top*) and Flag-BRCA1^Δ11^:6xHis-BARD1 (*Bottom*) for different nucleosome variants. Traces correspond to the titrations of MBP-BARD1^ARD-BRCTs^ (0.1 nM-4000 nM) or Flag- BRCA1^Δ11^:6xHis-BARD1 (0.1 nM-2000 nM) against 5’ FAM-labelled nucleosomes (at 20 nM each). Nucleosomes were either chemically modified with dimethyl-lysine analogues at H4 K20, and/or chemically ubiquitylated at H2A K15. Data are shown as average of three independent experiments +/-SEM. *K*_d_ values are summarised in Supplementary Table S1. **C.** EMSA experiments comparing MBP-BRCA1^RING^-BARD1 binding to H2AKc15ub and H2AKc15ub/H4Kc20me2 di-nucleosomes. Di-nucleosome variants were wrapped with 5’ FAM-labelled DNA, and increasing concentrations (8-512 nM) of MBP-BRCA1^RING^-BARD1 protein were incubated prior ot separation on native PAGE. Both di-nucleosomes were ubiquitylated at H2A position 15, and the H2AKc15ub/H4Kc20me2 variant was additional chemically modified with dimethyl-lysine analogues at H4 K20. Complexes were resolved by native-PAGE and imaged for fluorescein. **D.** Ubiquitylation assays using BRCA1^RING^-BARD1 and recombinant di-nucleosomes. Nucleosomes were either chemically ubiquitylated at H2A position 15 only (*Left*), modified with dimethyl-lysine analogues at H4 K20 and chemically ubiquitylated at H2A position 15 (*Middle*), or comprising H2AKc15ub-H4Kc20me2 and H2AKc15ub marks on different protomers (*Right*). Samples were taken prior to addition of E1 (-E1) and at 5, 15, 30 and 60 minute time points, and quenched by addition of 2x SDS loading buffer. Samples were resolved on SDS- PAGE gels and imaged for Alexa647 signal, before staining with Coomassie. Top panel shows the Alexa647 signal and lower panel a Coomassie stain of the total protein.

We examined how the presence of H4K20me2 would affect the binding and ubiquitylation activity of BRCA1-BARD1 *in vitro*. We produced recombinant mono- and di- nucleosomes with H4Kc20me2 and/or H2AKc15ub marks by chemical approaches, and tested BRCA1-BARD1 binding and activity in MST, EMSA and ubiquitylation assays. MST experiments showed a drastic reduction of BARD1^ARD-BRCTs^ binding to H4Kc20me2 mono- nucleosomes compared to unmodified nucleosomes (Figure 7B, *upper panel*, Supplementary Table S1), as observed previously in pull-down assays (33). Combination of H2A ubiquitylation and H4 methylation (H2AKc15ub/H4Kc20me2) marks partially rescued binding, although the *K*_d_ values were still much poorer than H2AKc15ub only nucleosomes (Figure 7B, *upper panel*, Supplementary Table S1). Similarly, BRCA1^Δ11^:BARD1 had reduced binding affinity for both H4Kc20me2 and H2AKc15ub/H4Kc20me2 nucleosomes (Figure 7B, *lower panel*,

Supplementary Table S1), and we observed comparable results for the BRCA1^RING^-BARD1 complex when assayed in the presence of the corresponding di-nucleosome variants in EMSA experiments (Figure 7C). Consistent with these data, ubiquitylation assays showed a reduction in BRCA1^RING^-BARD1 E3 ligase activity on H4Kc20me2 and H2AKc15ub/H4Kc20me2 nucleosomes compared to unmodified or H2AKc15ub substrates (Supplementary Figure S9A). These data show that H4K20me2 reduces the binding affinity of BARD1^ARD-BRCTs^ and BRCA1-BARD1 to both mono- and di-nucleosomes.

H4K20 methylation status is at its lowest during S/G2 phase of the cell cycle (90,91) due to dilution of parentally-marked histones after duplication of the genome. However, there is still significant H4K20me2 that has been inherited from parental chromatin, roughly equating to every other nucleosome containing this mark. These parental histones would block ARD interaction and presumably recruitment of BRCA1-BARD1 to chromatin, thus suggesting that only half of all available nucleosomes are conducive to BARD1^ARD-BRCTs^ binding. However, despite this inhibitory mark, BRCA1-BARD1 is still recruited and performs its function at DSB sites in S/G2.

This paradoxical observation can be reconciled based on the ‘bridge’ model (Figure 5A, Figure 7A). In this scenario BRCA1-BARD1 would preferentially span between adjacent nucleosomes, with the BARD1^ARD-BRCTs^ binding to the unmethylated newly-deposited nucleosome particle while the BRCA1^RING^-BARD1^RING^ heterodimer interacts with the adjacent di-methylated nucleosome (Figure 7A). To test this hypothesis we produced heterotypic alternating di-nucleosomes bearing unmodified H4 and H4Kc20me2 on adjacent nucleosome particles, both of which were also characterized by the H2AKc15ub mark (Supplementary Figure S9B, *Middle*), and tested this substrate together with H2AKc15ub (Supplementary Figure S9B, *Top*) and H2AKc15ub/H4Kc20me2 (Supplementary Figure S9B, *Bottom*) homotypic symmetric di-nucleosomes. The heterotypic di-nucleosome mixture showed an intermediate binding between the H2AKc15ub and H2AKc15ub/H4Kc20me2 symmetric substrates (Supplementary Figure S9C), supporting the ‘bridge’ model. To facilitate purification, and deconvolute H2A C-terminal tail ubiquitylation, we removed the 6xHis-tag present on H2A K15-linked ubiquitin from nucleosomes containing unmodified H4 while keeping the tag intact on H4Kc20me2 modified substrates (Supplementary Figure S9B and D). As such, RING-mediated H2A C-terminal ubiquitylation can be visualised as lower bands if taking place on the unmethylated nucleosomes. Conversely, ubiquitylation on H2AKc15ub- His/H4K20me2 marked protomers would run higher in the gel. We observed rapid ubiquitylation of H2AKc15ub symmetric di-nucleosomes (Figure 7D, *Left*) and lower activity on symmetric H2AKc15ub-His/H4K20me2 modified substrates (Figure 7D, *Middle*), likely due to reduced recruitment of BARD1^ARD-BRCTs^. In the presence of alternating hybrid modified di- nucleosomes, the H2AKc15ub-His/H4Kc20me2 nucleosome particle was ubiquitylated more rapidly compared to the corresponding symmetric substrate (Figure 7D, *Middle* and *Right,* Supplementary Figure S9E). These data suggest that BRCA1-BARD1 prefers bridging across nucleosomes, and can tolerate H4 di-methylation when this post-translational mark is adjacent to an unmethylated nucleosome particle. These observations provide an explanation for how BRCA1-BARD1 can bind to chromatin, and suggest that the biological-relevant substrate for BRCA1-BARD1 recruitment is a mother-daughter nucleosome.

## Discussion

Here we have shown that BRCA1-BARD1 combines multiple chromatin interaction domains to preferentially bridge across two nucleosomes simultaneously. Ubiquitin recognition is specific to the RNF8- and RNF168-induced H2AK13/15ub mark and can tolerate K63-linked ubiquitin chains at this position. BRCA1-BARD1 also recognises generic nucleosome features, including the acidic patch and H4 tail, providing both affinity and specificity. We have consistently seen that acidic patch mutations abrogate all binding and are the most deleterious variant in our assays. This is likely due to reliance on the acidic patch for both BARD1^ARD-BRCTs^ interaction with nucleosome (this work and Hu et al. (22)) and the RING domains interaction (21).

Interestingly, there is a difference in affinity for the two separable chromatin recognition regions. The RING domains alone bind nucleosomes with low affinity (21), while the BARD1^ARD-BRCTs^ integrate multiple signals to form a much larger interface with the nucleosome (Supplementary Figure 4A, RINGs-nucleosome interface = ∼1200 Å^2^, ARD-BRCTs- nucleosome interface = ∼3300 Å^2^). In the context of the BRCA1-BARD1 complex this affinity is additive suggesting both can engage concurrently, but we believe this is chiefly driven by the BARD1^ARD-BRCTs^ region of BARD1. Biologically, the specificity of the BARD1^ARD-BRCTs^ would ensure the BRCA1-BARD1 complex is recruited to the correct position in the genome, while the RING domains are then concomitantly to bind and ubiquitylate distal residues at the end of the H2A C-terminal tails. However, other important interactions can occur and a recent study has identified a novel DNA binding interface in the intrinsically disordered region between RING and ARD domains of BARD1 (35). This is consistent with our AFM observations showing an additional height in-between nucleosomes, presumably corresponding to BRCA1-BARD1 binding to linker DNA. In addition, we observed that a BRCA1-BARD1 region appears to initially dock to a nucleosome with high residency times while other interactions were more transient, suggesting a searching mechanism towards adjacent nucleosomes and dynamic association and dissociation of catalytic RING domains.

Bridging has been observed for a number of chromatin-interacting proteins (92–95) and this is critically important for their biological function. This mode of chromatin interaction for BRCA1-BARD1 may not be limited to cross-linking between adjacent nucleosomes on the same DNA strand. BARD1 contains an approximately 150 residue intrinsically disordered region separating the RING and ARD-BRCT domains allowing a great deal of flexibility in positioning of the two chromatin interacting regions. Indeed, our AFM and cryo-EM data suggest that, in isolation, the BRCA1-BARD1 complex is highly flexible. It is tempting to speculate that BRCA1-BARD1 may help HDR by being able to cross-link between sister DNA strands *in trans* as these would be in closer proximity after recent DNA synthesis in S-phase.

Interestingly, we could also visualize BRCA1-BARD1 bridging between mono- and di- nucleosome particles in AFM (Supplementary Figure S8D), suggesting the complex may also recognize non-connected nucleosomes as substrates.

The absence of the highly abundant H4K20me2 mark, and subsequent recognition of its unmodified state, enables differentiation between old and new chromatin. This occurs after dilution of parental histones during DNA replication, and guides complexes involved in DNA pathway choice (32,33,91). Given that BRCA1-BARD1 is unable to bind H4K20me2, and must outcompete its antagonist 53BP1 for damaged chromatin in S/G2, it was unclear as to how BRCA1-BARD1 could bind chromatin that is only marginally depleted of this mark. We suggest that this is accommodated by BRCA1-BARD1 utilizing two chromatin binding modules and bridging across nucleosomes. As such older H4K20me2 marked parental histones on can be bound by the tolerant RING domains and newly deposited histones by the ARD-BRCTs.

Outside of these two chromatin interaction regions BRCA1-BARD1 contains other well characterised domains, including those suggested to bind to DNA (35,75). We see little direct effect of the BRCA1-BRCT domains on chromatin interaction, a result that agrees well with cell biology-based observations (33,96). However, the BRCA1 BRCTs have been well characterised as phospho-protein interactors (25,27–29) that play a role downstream in DNA damage response signalling. Furthermore, the BRCA1 BRCT interactions are vital for response to more persistent breaks, as characterised through recruitment of RAP80 complexes many hours after damage induction (26), as well as recruitment to other DNA lesions such as replication stalls (97,98).

We surprisingly did not see any density for the RING domains in our structure on a mono-nucleosome (Figure 4B, Supplementary Figure S5B). However, subsequent biochemical analysis showed that clamping of BRCA1-BARD1 on mono-nucleosomes was possible, if not preferred (Figure 4D). It must be noted that absences of density in our cryo-EM map are not proof against this complex existing in solution, due to observation bias introduced during sample preparation (e.g. sample blotting, interaction with carbon, stability at air-water interface). Indeed, it has been shown previously that the RING domains are stabilised by the addition of E2 enzyme (21,22) not included in our preparations. Indeed, all the fragments of BRCA1-BARD1 can ubiquitylate mono-nucleosomes, albeit with lower activity. We suggest that while clamping is possible, BRCA1-BARD1 preferentially bridges between nucleosomes and spreads across chromatin. Further structural work to understand binding of the full BRCA1-BARD1 complex on chromatin would be required to expand on our observations.

In this study we have reconstituted ubiquitylation of H2A by the BRCA1-BARD1 RING domains in the context of larger BRCA1-BARD1 fragments. Despite extensive work (18,21,22), the exact role for H2A K125/127/129 ubiquitylation is not clear. While loss of ubiquitylation activity in mice has little effect on tumour development (99), human cells show increased cisplatin sensitivity (100). BRCA1 mediated ubiquitylation has been implicated in male meiosis (99), suppression of microsatellite repeat instability (19), and long-range DNA end resection (11). Here we show that the BARD1^ARD-BRCTs^ cannot bind to H2A K125/127/129ub, suggesting another reader may be implicated. Alternatively, ubiquitylation on the C-terminus of H2A is adjacent to entry/exit DNA and may play a direct role in affecting overall chromatin structure. It would be a fascinating avenue for future studies to characterise the molecular details of this ubiquitylation mark.

It was recently suggested that BARD1 ARD-BRCT binding blocks productive K63- linked ubiquitin chain elongation on H2A *in vitro* (22). We see that similar, if not better affinity for nucleosomes bearing pre-formed K63-linked di-ubiquitin at H2AK15, suggesting that if longer chains are assembled prior to BARD1 interaction, they do not prevent BRCA1-BARD1 recruitment. The actual nature of K63-linked chains in DDR remains unclear, with the RNF8 K63 target reported on multiple substrates (31,79,101). One component that is recruited to K63-linked ubiquitin chains at damage sites is RAP80, a mmember of the BRCA1-A complex that also includes BRCA1-BARD1. The presence of multiple ubiquitin binding domains on separate proteins within the BRCA1-A complex makes it interesting to understand if RAP80 could co-exist with BARD1 on the same ubiquitin chain or impart additional bridging interactions with other ubiquitylated proteins on chromatin. Indeed, the exact chain of events that leads to the recruitment of this complex and whether this a separable recruitment step to the BRCA1-BARD1 H2AK15ub di-nucleosome engagement described here would be a fascinating avenue of future studies.

### Data Availability

The cryo-EM map and associated structural model for the cat BARD1^ARD-BRCTs^:H2AKc15ub nucleosome complex have been deposited in the Electron Microscopy Data Bank and Protein Data Bank under accession codes EMD-16859 and 8OFF respectively. The cryo-EM maps for the isolated cat BRCA1^Δ11^:BARD1 complex can be accessed from the Electron Microscopy Data Bank under the accession codes EMD-16869 (Map 1, “closed” state) and EMD-16870 (Map 2, “open” state).

## Acknowledgments

We thank members of the Wilson and Zeqiraj labs for helpful discussions, and Roger Greenberg and Amélie Fradet-Turcotte for critical reading of the manuscript. We thank Daniel Durocher and Patrick Sung for gifts of BRCA1 and BARD1 plasmids. Human histone genes were sourced from Addgene and the Landry lab. We thank Logan Mackay in SIRCAMS, School of Chemistry, University of Edinburgh for 1D mass spec analysis.

## Author Contributions

M.F., E.Z. and M.D.W. conceived the study and supervised the project. H.B., M.F., E.Z. and M.D.W. designed the experiments (unless otherwise stated), analyzed the data, and wrote the manuscript, with input from the other authors. L.J.M., L.J.C., G.C. and D.K. purified protein and DNA components. Cryo-EM image processing and model building was performed by M.F. Biochemical assays were performed by H.B. and M.F. AFM experiments were performed and analysed by G.R.H.

