## Supplemental Files for "BRCA1-BARD1 combines multiple chromatin recognition modules to bridge nascent nucleosomes"

### Supplementary Figure Legends

**Supplementary Figure S1: Reconstitution of modified nucleosomes and quality control of assays.**

**A.** SDS-PAGE gel showing GST-BARD1$^{ARD-BRCTs}$ and MBP-BARD1$^{ARD-BRCTs}$ proteins used in pulldown, EMSA and MST assays. Gels were stained and imaged with colloidal Coomassie stain.

**B.** SDS-PAGE gel showing various ubiquitylated nucleosomes used in GST pulldown experiments. The gel was stained and imaged with colloidal Coomassie. Ub = Ubiquitin, AP = acidic patch.

**C.** Native-PAGE gel showing various ubiquitylated nucleosomes used in GST pulldown experiments. The gel was stained and imaged with Diamond DNA stain; shift in mobility suggests DNA is bound to histones. Doublet commonly observed for ubiquitylated nucleosomes.

**D.** SDS-PAGE gel representing the different steps required for chemical ubiquitylation of H2A mutants. Cysteine mutants are introduced at the desired site of ubiquitylation (e.g., H2AK15C), and to the terminal glycine of 6xHis-TEV-tagged Ubiquitin (Ub$^{G76C}$). H2AK15C and 6xHis-Ub$^{G76C}$ are mixed with di-bromo acetone (DBA), incubated at 4°C for ~30 minutes, and quenched with β-mercaptoethanol before being purified by ion exchange (IEX) and immobilised metal affinity (IMAC) chromatography. TEV protease is used to remove the 6xHis-tag from the ubiquitylated histone, and a second round of IMAC removes the protease and uncleaved samples. The gel was stained and imaged with colloidal Coomassie.

**E.** EMSA experiments assessing 6xHis-MBP-BARD1$^{ARD-BRCTs}$ protein binding to recombinant nucleosomes under increasing concentrations of NaCl. Ubiquitin specificity is increased at more physiological salt concentrations. Nucleosomes were wrapped with 5' FAM-labelled DNA and chemically ubiquitylated at H2A position 15. Complex was pre-formed with 2.3 nM nucleosome and 640 nM BARD1 in 100mM NaCl (seen to fully shift to complex band in both unmodified and H2AK15ub nucleosomes). Salt concentration was increased by 12.5 mM NaCl along the series from 112.5-200mM NaCl, showing loss of complex band for unmodified nucleosomes at salt concentrations. Complexes were resolved by native-PAGE and imaged for fluorescein.

**F.** SDS-PAGE gel showing various ubiquitylated nucleosomes used in EMSA, MST and GST pulldown experiments. The gel was stained and imaged with colloidal Coomassie.

**G.** Native-PAGE gel showing various nucleosomes used in EMSA and MST experiments. The gel was imaged for fluorescein. Ubiquitylated species run as multiple bands. Higher mobility of H2AKc15ub/H4Kc20me2 due to His tag on H2Aub.

**H.** Raw MST traces of 6xHis-MBP-BARD1$^{ARD\text{-}BRCTs}$ (*Left*) and Flag-BRCA1$^{\Delta 11}$:6xHis-BARD1 (*Right*) titrated against 5' FAM-labelled H2AKc15ub and acidic patch mutated nucleosomes. Refer to Figures 1F and 2B for details.

**Supplementary Figure S2: BRCA-BARD1 complexes and fusion proteins bind and ubiquitylate nucleosomes.**

**A.** SDS-PAGE gel showing 6xHis-MBP-BRCA1$^{RING}$-BARD1 Flag-BRCA1$^{\Delta 11}$:6xHis-BARD1, dStrepII-muGFP-BRCA1$^{\Delta 11}$:6xHis-BARD1 proteins used in assays. The gel was stained and imaged with colloidal Coomassie stain.

**B.** EMSA experiments using H2AKc15ub nucleosomes wrapped with 5' FAM-labelled DNA, and increasing concentrations (6.1-200 nM) of 6xHis-MBP-BRCA1$^{RING}$-BARD1 and dStrepII-muGFP-BRCA1$^{\Delta 11}$:6xHis-BARD1. Complexes were resolved by native-PAGE and imaged for fluorescein.

**C.** EMSA experiments using H2AKc15ub-AP nucleosomes wrapped with 5' FAM-labelled DNA, and increasing concentrations of (8-512 nM) of 6xHis-MBP-BRCA1$^{RING}$-BARD1. Complexes were resolved by native-PAGE and imaged for Fluorescein.

**D.** Schematic of the ubiquitylation assays performed using recombinant nucleosomes and BRCA1$^{RING}$-BARD1. Nucleosomes were combined in assay buffer with ATP, Ubiquitin (Ub), Ubiquitin$^{Alexa647}$ (Ub$^{Alexa647}$), UbcH5c (E2), and BRCA1$^{RING}$-BARD1 (E3). Reactions were initiated by addition of E1 enzyme.

**Supplementary Figure S3: Generating K63-linked ubiquitin chains on H2AKc15ub.**

**A.** SDS-PAGE gel showing the E1 activating and E2 conjugating enzymes used in E3 ligase activity assays or for the enzymatic ubiquitylation of histone H2A. The gel was stained and imaged with colloidal Coomassie.

**B.** SDS-PAGE gel showing a summary of the E2-mediated Lys-63 ubiquitylation reaction on chemically modified H2AKc15ub. Chain elongation beyond di-Ub was blocked using the K63R ubiquitin mutant. Additional bands indicated with asterisk correspond to by-products formed during the chemical ubiquitylation reaction, subsequently purified away in **C**. The gel was stained and imaged with colloidal Coomassie.

**C.** Purification process of H2AKc15ub-ub$^{K63R\text{-}HIS}$. Ion exchange (IEX) chromatography was used to remove unreacted ubiquitin and to partially separate the H2AKc15ub and H2AKc15ub-

ub$^{K63R}$ species. H2AKc15ub-ub$^{K63R}$ was further purified from H2AKc15ub by immobilised metal affinity chromatography (IMAC), taking advantage of a 6xHis-tag fused to ub$^{K63R}$. The tag was removed using TEV protease prior to wrapping the histone into octamers. Gels were stained and imaged with colloidal Coomassie.

**D.** SDS-PAGE gel showing H2AKc15ub and H2AKc15ub-ub$^{K63R}$ octamers. Gels were stained and imaged with colloidal Coomassie.

**E.** Native-PAGE gels showing unmodified, H2AKc15ub and H2AKc15ub-ub$^{K63R}$ nucleosomes. Ubiquitylated nucleosome run as doublets due to alternate conformations of covalently attached ubiquitin. Nucleosomes containing K63 chains are larger and have lower overall electrophoretic mobility. Gels were imaged for fluorescein.

**Supplementary Figure S4: Biochemical reagents probing the clamp hypothesis of BRCA1-BARD1 interaction.**

**A.** Cartoon model of mono-nucleosome core particles (from this study) showing overlapping interfaces of the two BRCA1-BARD1 Interaction modules. The H2A/H2B and H4 surfaces involved in BRCA1-BARD1 RINGs (*Left*) and BARD1 ARD-BRCTs (*Right*) interactions are coloured in light orange and green respectively. The histone octamer and nucleosomal DNA are coloured in light and dark grey.

**B**. Schematic of the method used for making asymmetrically-modified ubiquitylated octamers. The concentration of 6xHis-tagged ubiquitylated histone was limited in refolding leading to excess of unmodified octamer, which could be removed due to lack of a 6xHis-tag. Numbers next to individual histones indicate molar ratios of individual histone components mixed before dialysis into refolding buffer. Numbers next to folded octamers indicate the expected ratio of each species after refolding and after IMAC respectively. Octamers were subjected to SEC after IMAC.

**C.** SDS-PAGE gel showing unmodified, asymmetric and H2AKc15ub nucleosomes used in EMSA and ubiquitylation experiments. The gel was stained and imaged with colloidal Coomassie.

**D.** Native-PAGE gel showing unmodified, asymmetric and H2AKc15ub nucleosomes used in EMSA and MST experiments (*Left*). The gel was imaged for fluorescein. Nucleosomes were re-made and run for longer on native gel (*Right*) for better separation. Multiple bands observed for asymmetric nucleosome due to mixed species comprising mostly asymmetric modified, with minor unmodified and symmetrically modified nucleosomes each with multiple bands observed on native PAGE gel.

**Supplementary Figure S5: Cryo-EM structure determination and validation of BRCA1$^{\Delta11}$-BARD1:NCP$^{H2AKc15ub}$ in complex with H2Akc15ub nucleosomes.**

**A.** Representative micrographs (out of 16,015 movies collected) and 2D class averages (out of nine and 95 selected images following crYOLO and RELION particle picking, respectively) for the cryo-EM dataset of BRCA1$^{\Delta11}$:BARD1 in complex with H2AKc15ub nucleosomes. Red boxes and green circles (240 pixels each) indicate picked particles.

**B.** Flow-chart of data processing strategy. Boxed 3D initial models and refined class were selected for subsequent processing. The final map at a global resolution of 3.40 Å is shown at different thresholds and coloured by local resolution. Refer to Supplementary Table S2 for details.

**C.** FSC curves (*Left*) and Euler angle distribution (*Right*) in the final 3D reconstruction of the BARD1$^{ARD-BRCTs}$:NCP$^{H2AKc15ub}$ complex. The resolution was calculated using the gold-standard FSC cut-off at 0.143 frequency; rod heights are proportional to the number of particles in each direction.

**D.** Plot of the directional FSC (3DFSC) that represents a measure of directional resolution anisotropy. Global FSC (red lines) at a resolution of 3.40 Å, the spread of directional resolution values ± 1 standard deviation from the mean (area within the green dotted lines), and a histogram of 100 directional resolutions evenly sampled over the 3DFSC (blue bars) are indicated. A sphericity of 0.882 was determined at the gold-standard FSC cut-off of 0.143 frequency, indicating no significant anisotropic angular distributions of particles.

**E.** Overlays between the cryo-EM structures of BARD1$^{ARD-BRCTs}$:NCP$^{H2AKc15ub}$. The structure reported in this study is coloured as in **Figure 4B**, while the corresponding complexes reported previously (PDB ID: 7LYC; PDB ID: 7E8I) are coloured grey. Structures are shown in cartoon models.

**Supplementary Figure S6: Generation of di-nucleosomes for assaying BRCA1-BARD1 binding.**

**A.** EMSA experiments using various di-nucleosomes to assess the role of the acidic patch in binding. Di-nucleosomes wrapped with 5’ FAM-labelled DNA, and increasing concentrations (10-100 nM) of BRCA1$^{RING}$-BARD1. Complexes were resolved by native-PAGE and imaged for fluorescein.

**B.** Ubiquitylation assays assessing 6xHis-MBP-BRCA1$^{RING}$-BARD1 E3 ligase activity on acidic patch mutant (*Left*) and acidic patch mutant/H2AKc15ub (*Right*) di-nucleosomes. Samples were taken prior to addition of E1 (-E1) and at 5, 15, 30 and 60 minute time points, and quenched by addition of 2x SDS loading buffer. Samples were resolved on SDS-PAGE gels and imaged for Alexa647 signal, before staining with Coomassie. Top panel shows the Alexa647 signal, and lower panel a colloidal Coomassie stain of the total protein.

**C.** Model of heterotypic di-nucleosomes generated by mixing equimolar ratios of H2AKc15ub-His octamer and unmodified octamer in di-nucleosome wrapping, prior to IMAC based purification.

**D.** Native-PAGE gel showing the wrapping of unmodified, H2AKc15ub and unmodified/H2AKc15ub heterotypic di-nucleosomes. Multiple bands expected due to presence of doublet-forming ubiquitylated species with and without a 6xHistag which affects mobility on native PAGE.

**E**. Native-PAGE illustrating purification protocol used to enrich for heterotypic di-nucleosomes from **D**. Mixed population of fully ubiquitylated, partially ubiquitylated (only single octamer) and non-ubiquitylated di-nucleosome samples were incubated with Ni-NTA beads for two hours and then washed with IMAC A buffers, before being eluting from the beads with increasing concentrations of imidazole (*Right*). Recovered di-nucleosomes were buffer exchanged into nucleosome buffer and used in ubiquitylation assays.

**F.** Ubiquitylation assays testing 6xHis-MBP-BRCA1$^{RING}$-BARD1 ligase activity on the unbound fraction enriched for a species with a single nucleosome containing ubiquitin from heterotypic di-nucleosome purification in **E**. Samples were taken prior to addition of E1 (-E1) and at 5, 15, 30 and 60 minute time points, and quenched by addition of 2x SDS loading buffer. Samples were resolved on SDS-PAGE gels and imaged for Alexa647 signal, before staining with Coomassie. Top panel shows the Alexa647 signal, and lower panel a colloidal Coomassie stain of the total protein.

**Supplementary Figure S7: Cryo-EM structure determination and validation of BRCA1$^{\Delta 11}$:BARD1 alone data.**

**A.** Representative micrographs (out of 227 movies collected) and 2D class averages (out of 91 selected images) for the isolated BRCA1$^{\Delta 11}$:BARD1 complex dataset. Red boxes (220 pixels) indicate picked particles.

**B.** Flow-chart of data processing strategy. Boxed 3D initial models were selected for subsequent processing. The final maps at the reported global resolutions of 7.44 Å (*Left*; “closed” state) and 5.56 Å (*Right*; “open” state) are shown and coloured by local resolution. Refer to Supplementary Table S2 for details.

**C.** FSC curves (*Left*) and Euler angle distribution (*Right*) in the final 3D reconstruction of the BRCA1$^{\Delta 11}$-BARD1 “closed” state. The resolution was calculated using the gold-standard FSC cut-off at 0.143 frequency; rod heights are proportional to the number of particles in each direction.

**D.** As in **C**, but for the BRCA1$^{\Delta 11}$:BARD1 “open” state.

**Supplementary Figure S8: AFM imaging of di-nucleosomes and BRCA1$^{\Delta 11}$:BARD1 complex.**

**A.** AFM simulation of unmodified di-nucleosomes (PDB ID: 7PF4) and of the “open” state BRCA1$^{\Delta 11}$:BARD1 complex. Tip radius and sampling values used in simulation AFM were 1 nm and 3 pixels/nm, respectively. Refer to Figure 6D for details.

**B.** HS-AFM of di-nucleosome particles (*Left*) and close-up views captured over time (*Right*). Zoomed-in views were taken from a different area than the overview image.

**C.** HS-AFM of BRCA1$^{\Delta 11}$:BARD1:di-nucleosome complex imaged over time.

**D.** Overview images obtained by HS-AFM of BRCA1$^{\Delta 11}$:BARD1 interactions with unmodified di-nucleosomes over time. Red arrows indicated BRCA1$^{\Delta 11}$:BARD1 bridging between a mono- and a di-nucleosome; white arrows indicate BRCA1$^{\Delta 11}$:BARD1 bridging across a di-nucleosome.

**Supplementary Figure S9: Formation and testing H4 methylated nucleosomes and di-nucleosomes.**

**A.** Ubiquitylation assays assessing BRCA1$^{RING}$-BARD1 E3 ligase activity on H4Kc20me2 and H2AKc15ub/H4Kc20me2 mono- or di-nucleosomes. Samples were taken prior to addition of E1 (-E1) and at 5, 15, 30 and 60 minute time points, and quenched by addition of 2x SDS loading buffer. Samples were resolved on SDS-PAGE and imaged for Alexa647 signal, before staining with Coomassie. Top panel shows the Alexa647 signal and lower panel a colloidal Coomassie stain of the total protein.

**B.** Schematic representation of different di-nucleosomes generated to assess the role of H4K20me2 in a di-nucleosome context.

**C.** EMSA experiments using various di-nucleosomes wrapped with 5’ FAM-labelled DNA, and increasing concentrations (8-512 nM) of 6xHis-MBP-BRCA1$^{RING}$-BARD1. Complexes were resolved by native-PAGE and imaged for fluorescein.

**D.** SDS-PAGE (*Left*) and native (*Right*) gels showing the purification process of H2AKc15ub, H4Kc20me2 and H2AKc15ub-H4Kc20me2 heterotypic di-nucleosomes. Heterotypic nucleosomes were incubated with Ni-NTA beads for two hours, and then washed with IMAC A buffers, before eluting from the beads with increasing concentrations of imidazole.

Recovered di-nucleosomes were buffer exchanged into nucleosome buffer, then used in ubiquitylation assays.

**E.** Ubiquitylation assays testing BRCA1$^{RING}$-BARD1 activity on the unbound fraction from heterotypic di-nucleosome purification in **D**. Samples were taken prior to addition of E1 (-E1), and at 5, 15, and 30 minute time points, and quenched by addition of 2x SDS loading buffer. Samples were resolved on SDS-PAGE gels and imaged for Alexa647 signal, before staining with Coomassie. Top panel shows the Alexa647 signal, and lower panel a Coomassie stain of the total protein.

### Supplementary Figure S1

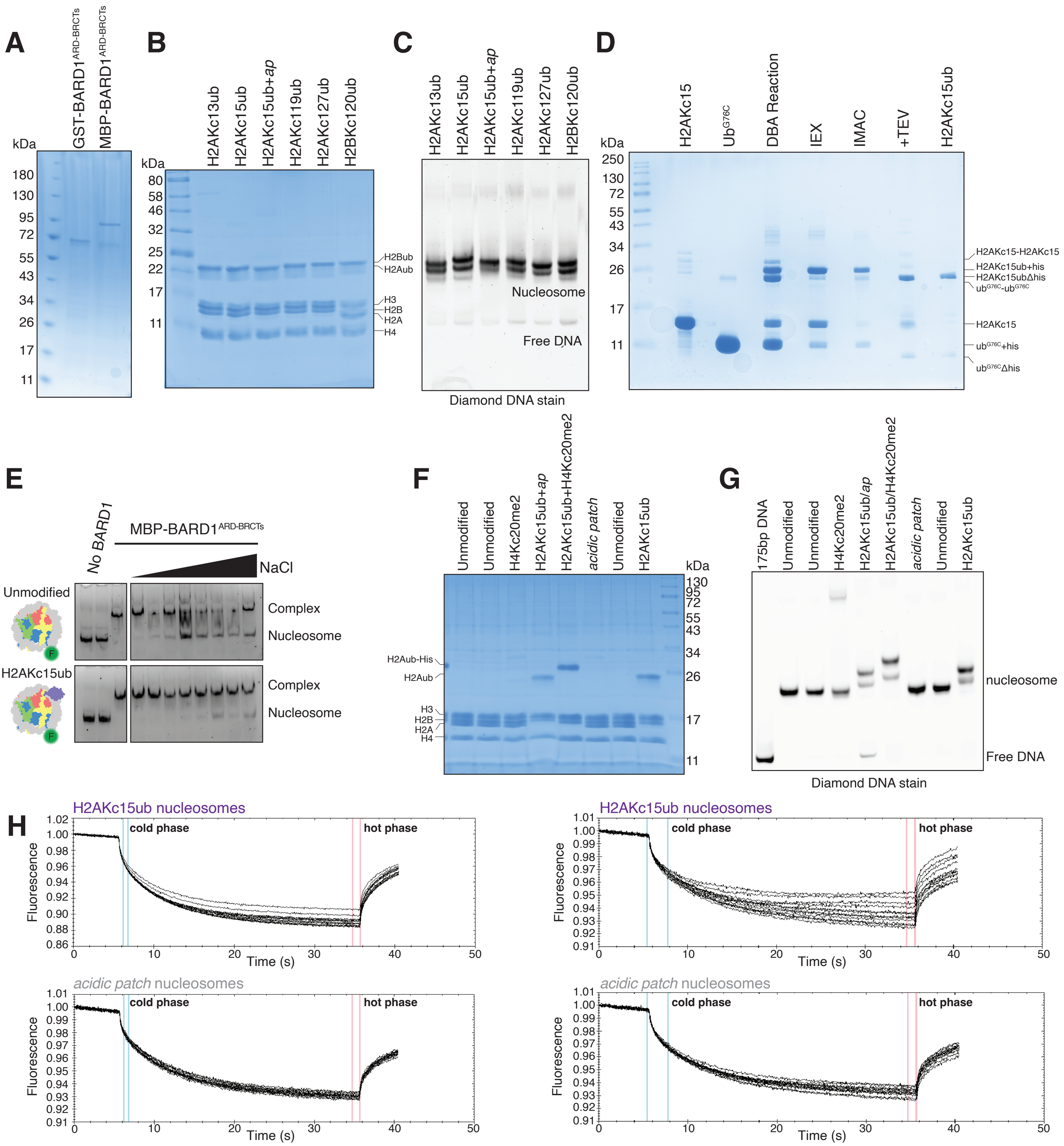

### Supplemetary Figure S2

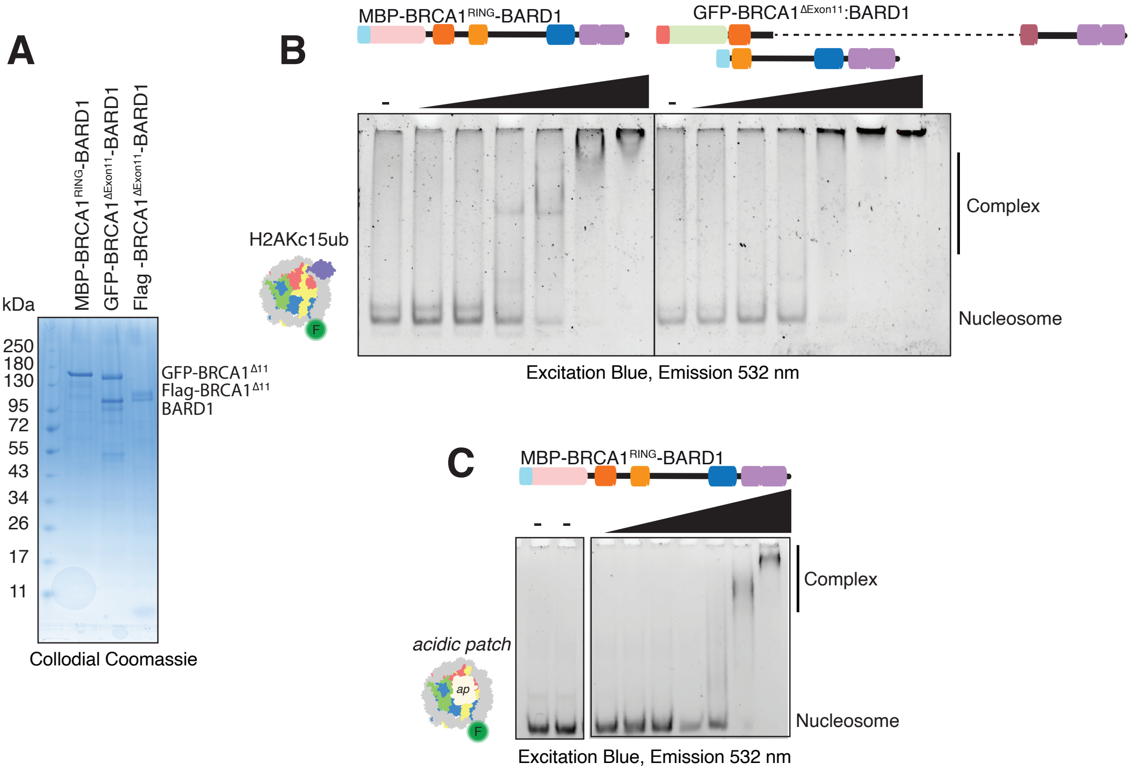


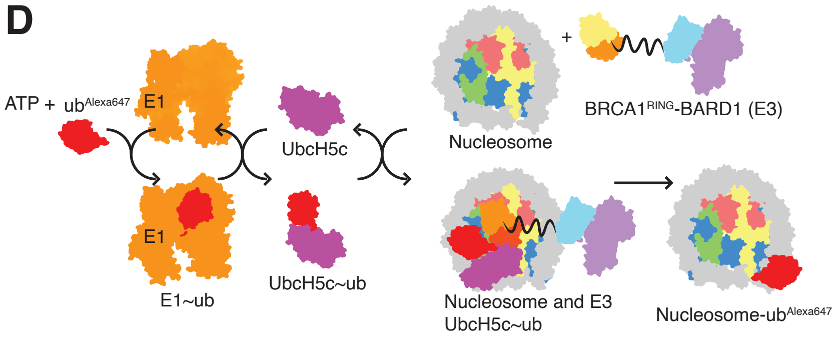

### Supplementary Figure S3

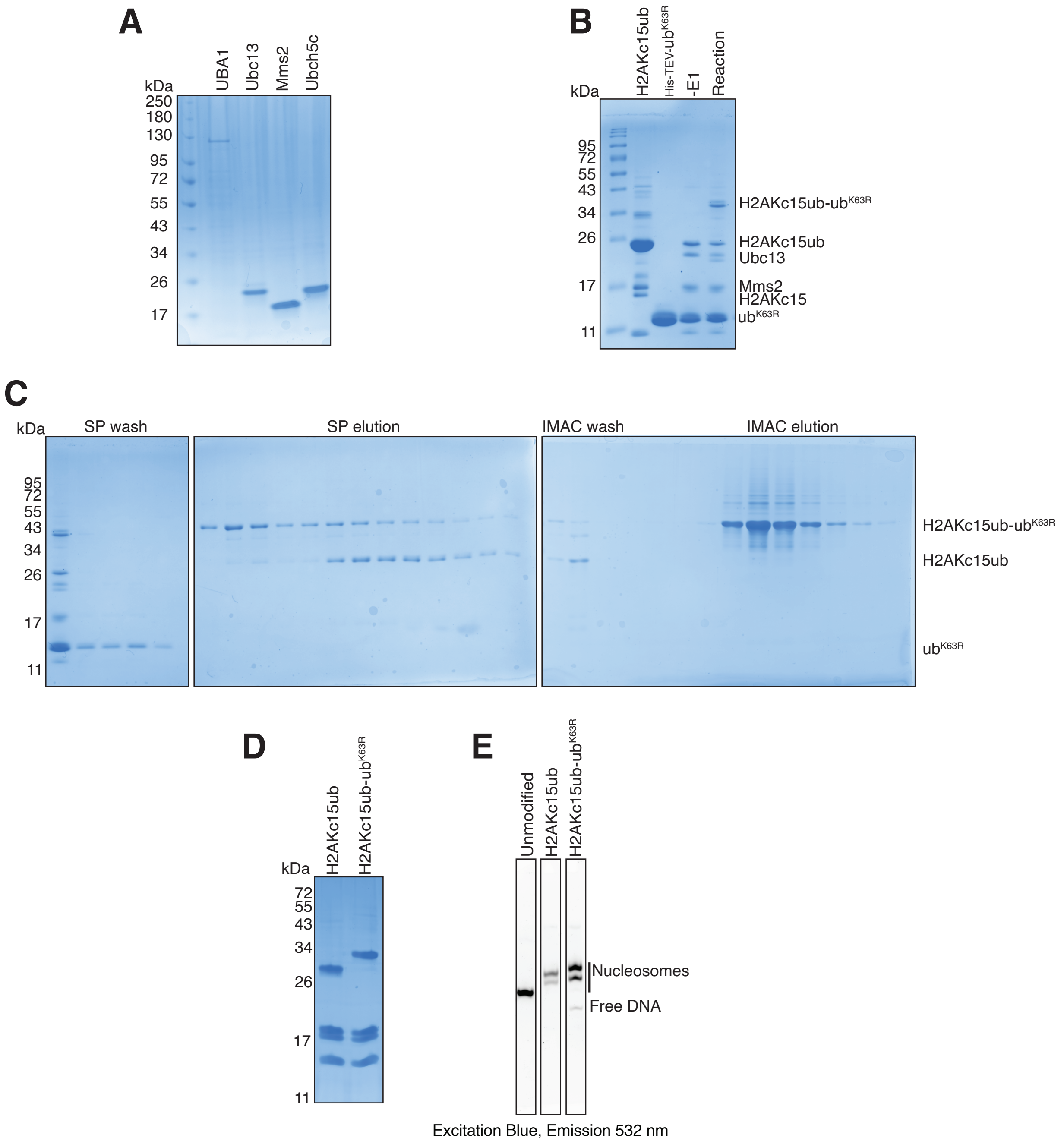

### Supplementary Figure S4

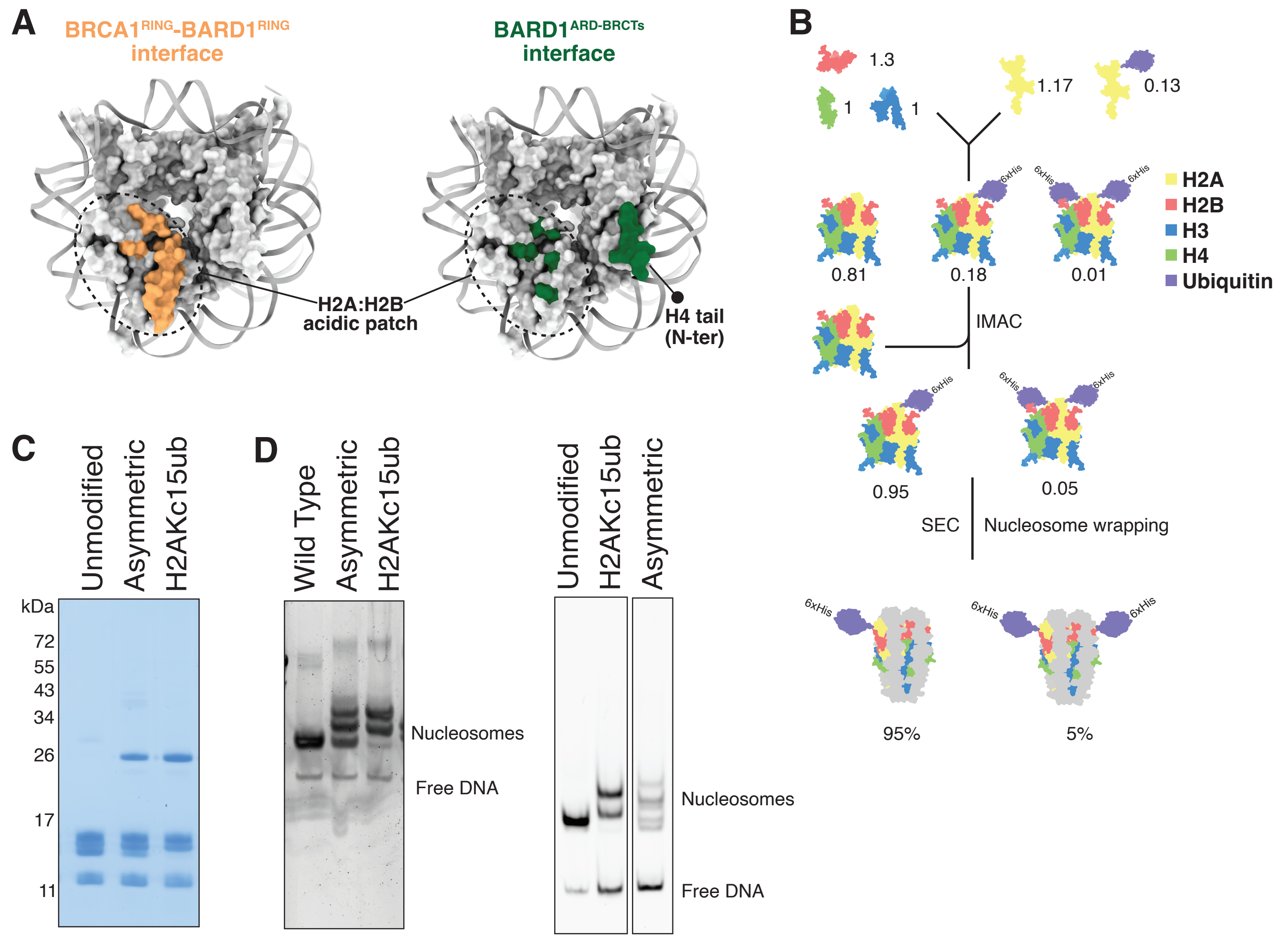

### Supplementary Figure S5

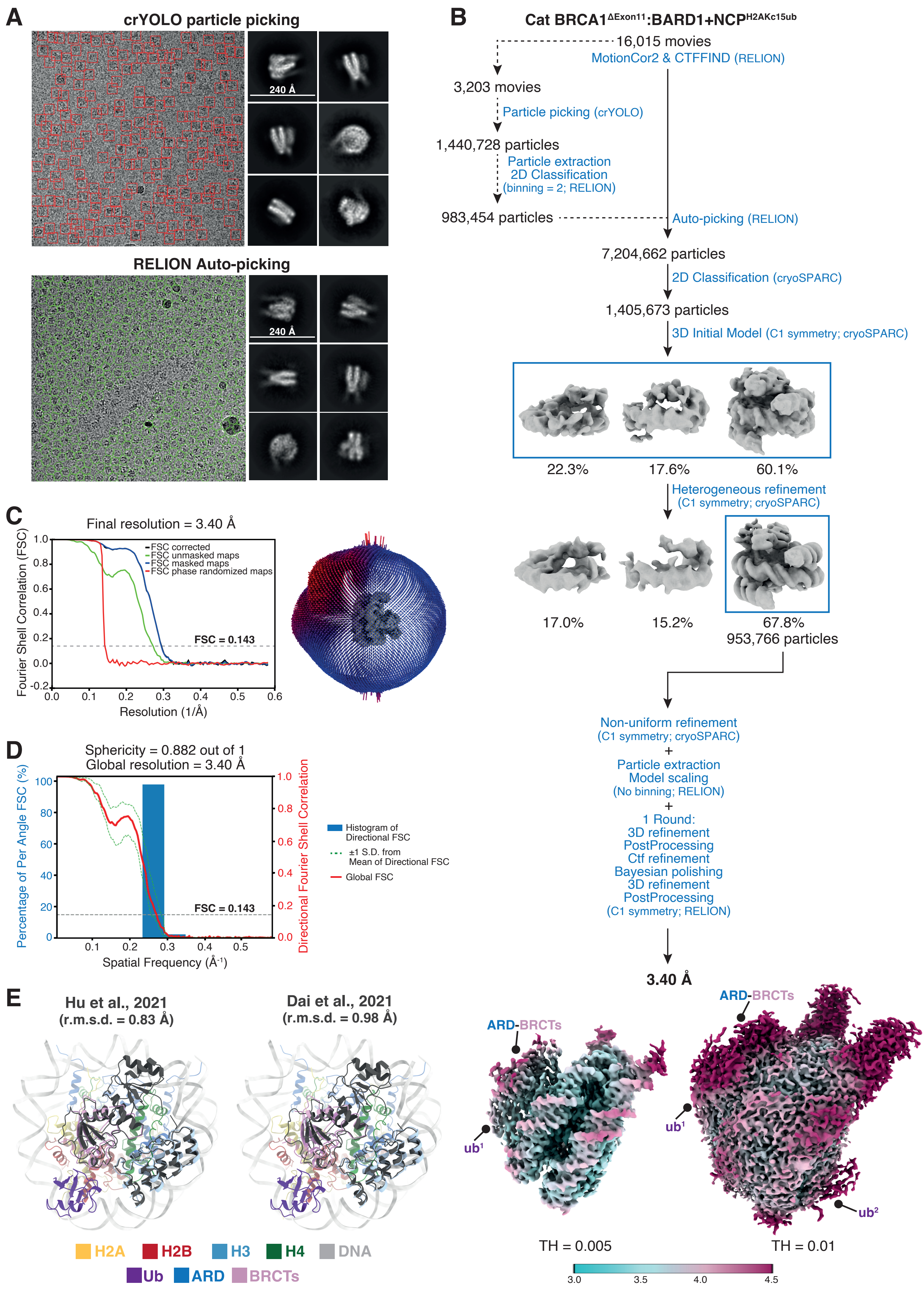

### Supplementary Figure S6

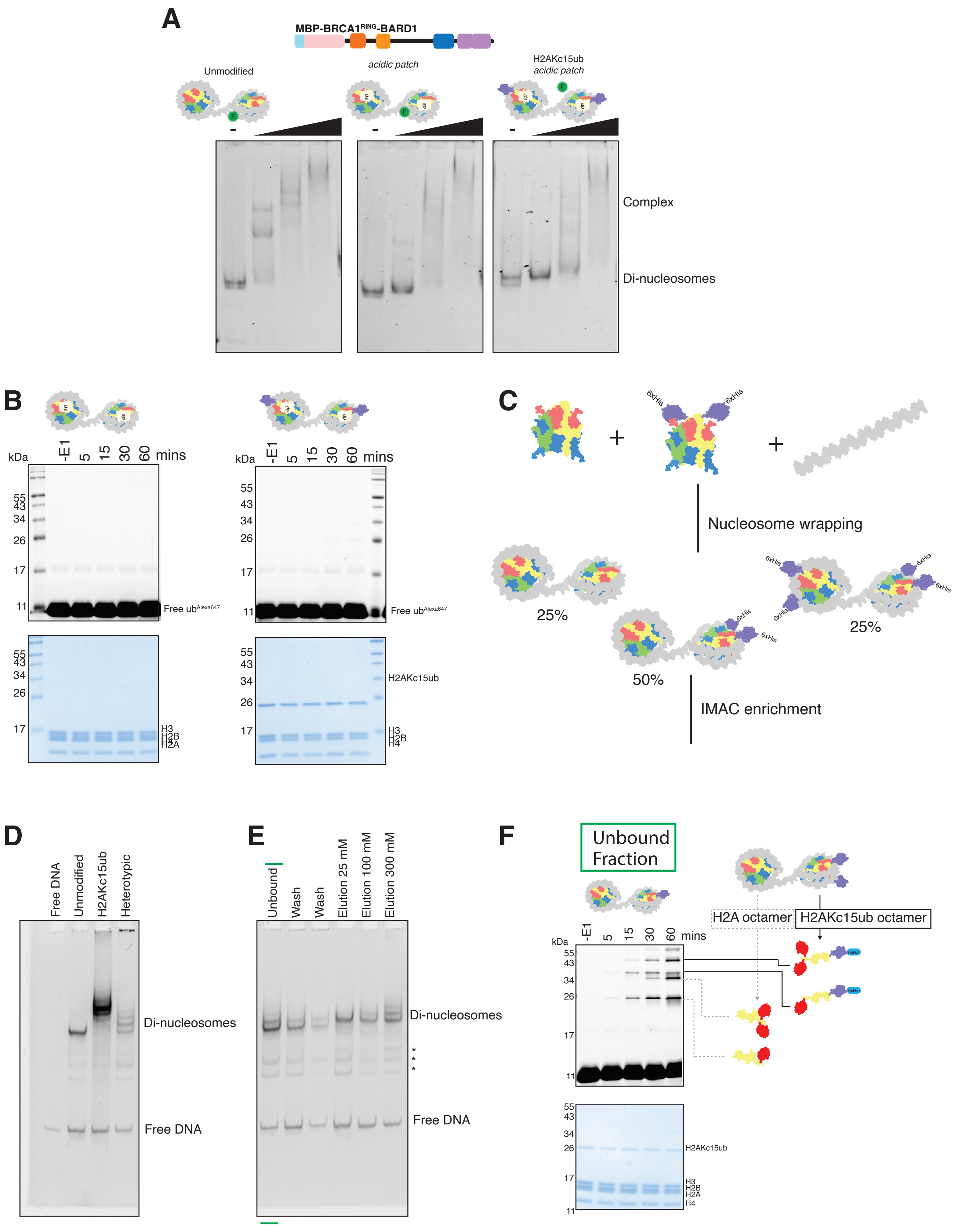

### Supplementary Figure S7

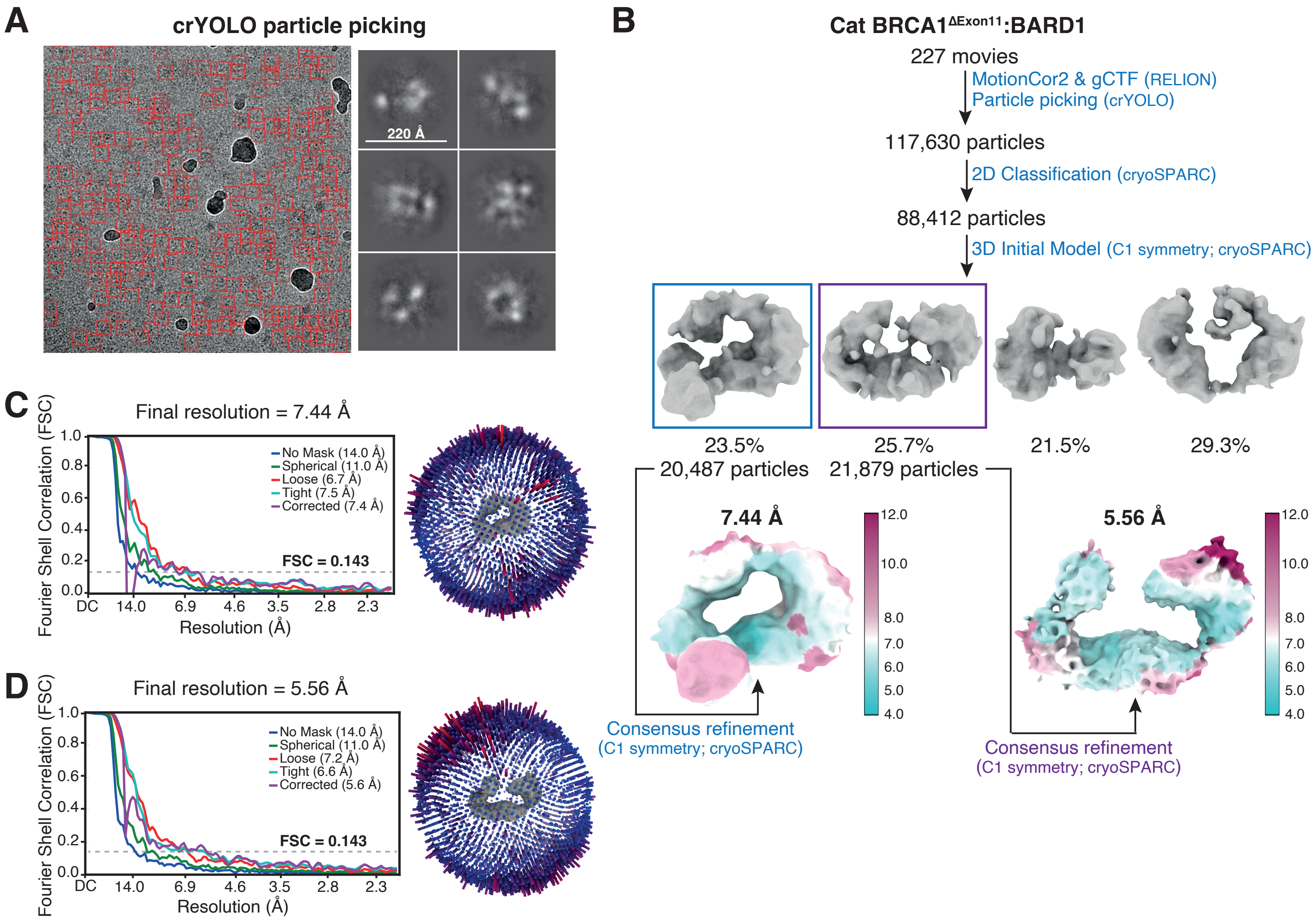

### Supplementary Figure S8

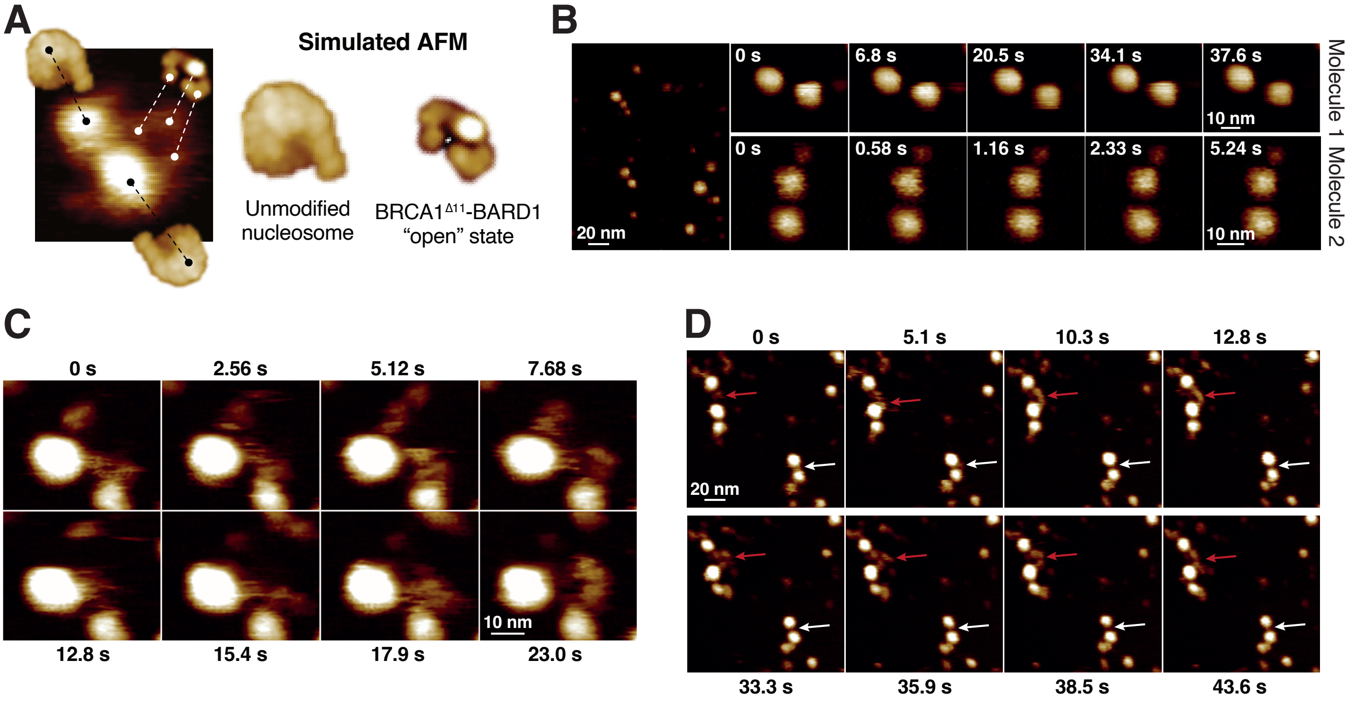

### Supplementary Figure S9

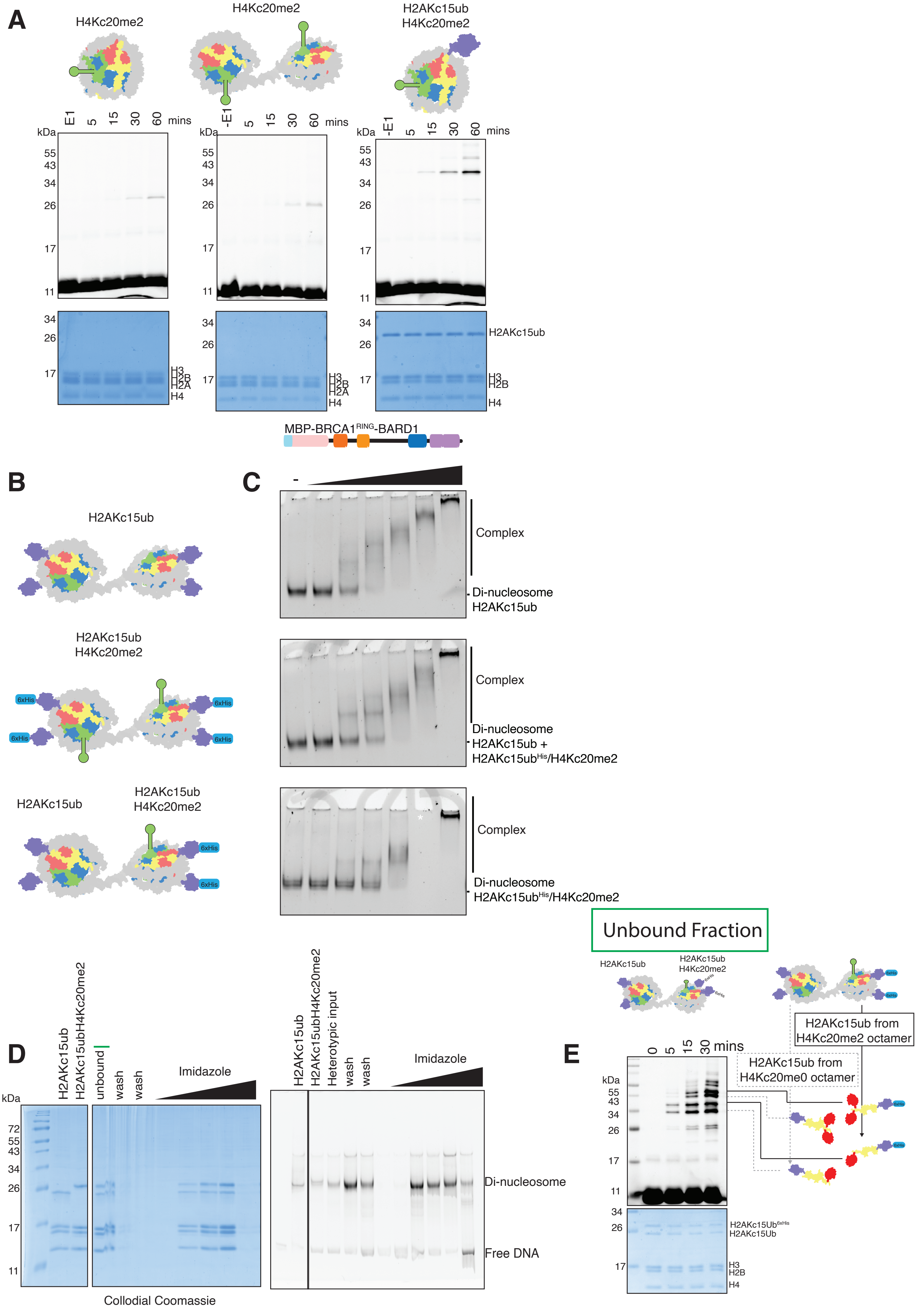

#### Supplementary Tables

##### Supplementary Table S1 – Summary of dissociation constant values ($K_d$) calculated by MST

| | $BARD1^{ARD-BRCTs}$ | $BRCA1^{\Delta 11:}BARD1$ |
|---|---|---|
| **Nucleosome variants** | | |
| Wild type | 0.218 μM ± 0.044 μM | 0.093 μM ± 0.026 μM |
| H2AKc15ub | 0.072 μM ± 0.010 μM | 0.034 μM ± 0.005 μM |
| H2AKc15ub-ub | 0.024 μM ± 0.004 μM | 0.013 μM ± 0.002 μM |
| H4Kc20me2 | 0.964 μM ± 0.889 μM | 0.365 μM ± 0.302 μM |
| Acidic Patch | ND | ND |
| H2AKc15ub + Acidic Patch | ND | ND |
| H2AKc15ub + H4Kc20me2 | 0.298 μM ± 0.157 μM | 0.118 μM ± 0.055 μM |

#### Supplementary Table S2 – Cryo-EM data collection, refinement and validation statistics

| | $BARD1^{ARD\text{-}BRCTs}$:$NCP^{H2AKc15ub}$ (EMDB-16859; PDB ID: 8OFF) | $BRCA1^{\Delta 11}$:BARD1 (EMDB-16869; EMDB-16870) | |
|---|---|---|---|
| **Data Collection** | | | |
| Microscope | TFS Titan KRIOS | TFS Titan KRIOS | |
| Detector | Falcon IVi | Falcon III | |
| Voltage (keV) | 300 | 300 | |
| Mode | Counting | Integrating | |
| Pixel size (Å) | 0.86 | 1.065 | |
| Magnification (x) | 96,000 | 75,000 | |
| Electron dose ($e^-$/Å$^2$) | 36.4 | 73 | |
| Exposure (s) | 5 | 1.7 | |
| EPU Frames (Nr) | 172 | 50 | |
| Electron dose per frame ($e^-$/Å$^2$) | 0.8 | 1.46 | |
| EER Fractions | 45 | - | |
| No. of movies | 16,015 | 227 | |
| Defocus range ($\mu$m) | -1.7 to -3.1 | -1.5 to -3.0 | |
| **Data Processing** | | **Map 1** | **Map2** |
| Symmetry Point Group | C1 | C1 | C1 |
| Initial particle number | 7,204,662 | 117,630 | 117,630 |
| Final particle number | 953,766 | 20,487 | 21,879 |
| Map resolution (Å) | 3.40 | 7.44 | 5.56 |
| FSC threshold | 0.143 | 0.143 | 0.143 |
| Map resolution range (Å) | 3.0-4.5 | 4.0-12.0 | 4.0-12.0 |
| **Refinement** | | | |
| Model resolution (Å) | 3.45 | | |
| FSC threshold | 0.5 | | |
| Map sharpening *B* factor (Å$^2$) | -150.71 | | |
| Model composition | | | |
| Chains | 16 | | |
| Non-hydrogen atoms | 13,922 | | |
| Protein residues | 1,057 | | |
| Nucleotide | 284 | | |
| Water | 0 | | |
| Ligands | 0 | | |
| *B* factors (Å$^2$) | | | |
| Protein (min/max/mean) | 7.18/73.86/29.94 | | |
| Nucleotide (min/max/mean) | 31.44/140.43/83.22 | | |
| Water | - | | |
| Ligands | - | | |
| Map:model CC | | | |
| CC (mask) | 0.86 | | |
| CC (box) | 0.79 | | |
| CC (peaks) | 0.74 | | |
| CC (volume) | 0.82 | | |
| R.m.s. deviations | | | |
| Bond lengths (Å) | 0.010 | | |
| Bond angles (°) | 0.801 | | |
| **Validation** | | | |
| MolProbity score | 2.48 | | |
| Clashscore | 5.01 | | |
| Rotamers outliers (%) | 9.17 | | |
| Cß outliers (%) | 0 | | |
| Peptide plane | | | |
| Cis proline/general (%) | 0/0 | | |
| Twisted proline/general (%) | 0/0 | | |
| CaBLAM outliers (%) | 2.52 | | |
| Ramachandran plot | | | |
| Favored (%) | 92.02 | | |
| Allowed (%) | 7.98 | | |
| Outliers (%) | 0 | | |

**Supplementary Table S3 – Summary of the BRCA1 and BARD1 expression constructs used in this study**

| Expression Construct | Species | Protein 1 (name) | Protein 1 (aa) | Tag | Protein 2 (name) | Protein 2 (aa) | Tag | Expression System |
|---|---|---|---|---|---|---|---|---|
| BARD1 ARD-BRCTs | Human | BARD1 | 425-777 | N-ter. GST | - | - | - | *E. coli* |
| BARD1 ARD-BRCTs | Human | BARD1 | 425-777 | N-ter. 6xHis-MBP | - | - | - | *E. coli* |
| BRCA1$^{RING}$-BARD1 | Human | BRCA1$^{RING}$ | 1-100 | N-ter. 6xHis-MBP | BARD1 | 26-777 | none | *E. coli* |
| BRCA1$^{\Delta 11:}$ BARD1 | Human | BRCA1$^{\Delta 11}$ | 1-223, 1366-1863 | N-ter. Flag | BARD1 | 1-777 | N-ter. 6xHis | *Tni* |
| BRCA1$^{\Delta 11:}$ BARD1 | Human | BRCA1$^{\Delta 11}$ | 1-223, 1366-1863 | N-ter. dStrepII-muGFP | BARD1 | 1-777 | N-ter. 6xHis | *Tni* |
| BRCA1$^{\Delta 11:}$ BARD1 | Cat | BRCA1$^{\Delta 11}$ | 1-223, 1369-1873 | N-ter. Flag | BARD1 | 1-773 | N-ter. 6xHis | *Tni* |
| BRCA1$^{\Delta 11:}$ BARD1 | Cat | BRCA1$^{\Delta 11}$ | 1-223, 1369-1873 | N-ter. dStrepII-muGFP | BARD1 | 1-773 | N-ter. 6xHis | *Tni* |
